# Cycling of sulfur redox intermediates drives microbial activity in the sulfate–methane transition zone of cold methane seeps

**DOI:** 10.64898/2025.11.30.691342

**Authors:** Eryn M. Eitel, Ranjani Murali, Daniel R. Utter, Fabai Wu, Sujung Lim, Stephanie A. Connon, Alex Sessions, Victoria J. Orphan

## Abstract

Microbial sulfate reduction is a cornerstone of marine sediment biogeochemistry, driving carbon remineralization and fueling the anaerobic oxidation of methane (AOM). Yet in zones of high methane flux, sulfate limitations may constrain the sulfate-reducing bacteria (SRB) and anaerobic methanotrophic archaea (ANME) that typically perform AOM. Although often overlooked, sulfur redox intermediates are readily utilized by diverse microorganisms, potentially driving AOM in sulfate-limited zones. To resolve the microbial mechanisms underlying cryptic sulfur cycling in such sediments, Monterey Canyon cold methane seeps were investigated through an integrated geochemical, isotopic, and metatranscriptomic approach. High-resolution electrochemical measurements confirmed intense sulfide production in seep sites, and long-term anoxic incubations were conducted with sediment from the SMTZ amended with elemental sulfur, thiosulfate, or sulfate as the sole sulfur source, with or without methane. Over 650 days, sulfide accumulation was greatest in elemental sulfur treatments, followed by thiosulfate and sulfate; in all cases methane addition enhanced sulfide production. Isotopic measurements showed modest S-isotope fractionation indicating that the large fractionations typical of slow sulfate reduction were muted by additional sulfur transformations. In elemental sulfur treatments, isotopic and geochemical patterns suggested that disproportionation was unlikely. Metatranscriptomes revealed broad expression of *sox* genes and abundant *dsr/apr* across treatments, along with thiosulfate-linked *phs* upregulation. While ANME-2c and SEEP-SRB2 activity increased with methane, transcriptomic and isotopic data together highlighted the roles of *Desulfocapsaceae, Desulfobulbaceae*, and *Sulfurovaceae* lineages in mediating sulfur transformations. Taken together, these results demonstrate how cryptic sulfur cycling may sustain microbial communities in sulfate-depleted deep-sea sediments and contribute to AOM.

## Introduction

Cold seeps are globally significant biogeochemical hotspots that fuel chemosynthetic primary production and host tightly coupled redox cycles. A hallmark process in these settings is the anaerobic oxidation of methane (AOM), often facilitated by direct interspecies electron transfer (DIET) within syntrophic consortia of anaerobic methanotrophic archaea (ANME) and sulfate-reducing bacteria (SRB) [1–4]. Through this partnership, the carbon and sulfur cycles are intimately linked, attenuating benthic methane fluxes [5]. Multicellular ANME-SRB consortia are particularly prevalent in cold methane seeps due to the rapid advection of methane-rich porewaters and prevalence of sulfate [6–9]. Paradoxically, the highest rates of AOM often occur within the sulfate-methane transition zone (SMTZ) where sulfate becomes depleted and diffusion alone may not be able to re-supply the concentrations required [10–13]. In these sulfate-limited sediments, cryptic sulfur cycling may sustain microbial communities and mediate critical biogeochemical transformations, including AOM.

In deep-sea sediments, particularly within the SMTZ, sulfur redox intermediates such as elemental sulfur, polysulfides, sulfite, and thiosulfate, may play important roles in fueling microbial processes. Although the concentrations of sulfur redox intermediates are typically low relative to sulfate, this is likely not representative of their importance [14]. In fact, low concentrations may result from rapid turnover [15–17], demonstrating the favorable microbial consumption of these sulfur species. In the low sulfate conditions of early Earth, sulfite and thiosulfate likely played prominent roles [18], and in modern environments the reduction or disproportionation of these compounds by the classically termed ‘sulfate-reducing’ bacteria is common [19–25].

While sulfate-dependent AOM has long dominated our understanding of methane seeps, new findings suggest a more complex network of sulfur transformations at play. In low sulfate settings resembling the SMTZ, AOM has been linked to sulfide oxidation products [13, 26, 27] and isotopic signals in seep sediments have been attributed to sulfide oxidation [28–32]. However, rather than regenerating sulfate, the biological and chemical oxidation of sulfide typically produces a range of sulfur redox intermediates. [33–36]. Although the role of these intermediates within methane seeps is not fully understood, elemental sulfur may indicate former SMTZ locations [37, 38] and polysulfide concentrations correlate well with rates of AOM [39]. Indeed, polysulfide-driven AOM has been proposed [40] and thiosulfate, sulfite, and elemental sulfur additions stimulated AOM in non-seep sediments [41–44].

Although sulfur redox intermediates may play an important role within the SMTZ, the mechanisms underlying their transformations remain poorly characterized. Cryptic sulfur cycling may sustain AOM under sulfate-limiting conditions, however the microbial communities and genomic pathways mediating these reactions are still unresolved. Additionally, the sulfur isotope fractionation associated with cycling of these sulfur redox intermediates is not well constrained. To identify the key microbial players and the processes connected to sulfur cycling within the SMTZ, long-term, anoxic microcosm incubations were established to test the response of marine sediment communities to different sulfur compounds. In these incubations, sulfate, thiosulfate, and elemental sulfur were independently added as the sole sulfur source to sediment collected from a cold seep in the submarine Monterey Canyon. To test the metabolic flexibility of the ANME-SRB partnership and the ability of these sulfur compounds to facilitate AOM, cycling of sulfur redox intermediates was examined both with and without methane. Over the course of 650 days, sulfur concentrations, sulfide isotope compositions, and microbial community dynamics were tracked. Additionally, metatranscriptomics was used as an endpoint measurement to assess differences in expression between treatments while illuminating the role of important taxa. The results demonstrate active and complex sulfur cycling between diverse phyla within a methane seep associated SMTZ and provide insight into sulfur isotope fractionation with implications for interpreting the geological record.

## Methods

### Sediment collection and shipboard processing

Sediment cores were collected from methane cold seeps in the Extrovert Cliff area of Monterey Canyon at a depth of approx. 960 m on the *R/V Western Flyer* and ROV *Doc Ricketts* in May 2018 and August 2019 (Table S1). Seeps previously identified within this area are known to contain high levels of sulfide and methane and all seep sites in this study were identified by visible fluid flow, yellow or peach-colored microbial mats, and encircled by macrofauna, such as vesicomyid clams [45–48]. A swing-arm apparatus mounted on the ROV *Doc Ricketts* enabled collection of long (1 m) sediment push cores in addition to standard length (30 cm) cores, which were either profiled with voltammetric microelectrodes, or extruded, sectioned, and processed shipboard for porewater geochemistry and microbiological analyses within 1 hour of retrieval.

High-resolution depth profiles of dissolved oxygen and sulfide were obtained in ex-situ cores with microelectrodes as previously described [49]. Briefly, Au solid state electrodes were constructed in glass housing and plated with Hg [50, 51]. The three-electrode configuration, made up of the Hg/Au voltammetric working microelectrode, an Ag/AgCl reference electrode, and a Pt counter electrode were controlled using a DLK-70 potentiostat with DLK MAN-1 micromanipulator (Analytical Instrument Systems Inc; NJ, USA). A combination of linear sweep and square-wave voltammetry was used to determine dissolved oxygen and sulfide [52] and data was integrated using VOLTINT, a semiautomated Matlab® script with peak recognition [53].

Cores processed for porewaters were sectioned in 1-3 cm thick horizons and porewaters were extracted using a pneumatic sediment squeezer (KC Denmark A/S; Silkeborg, Denmark) under N_2_ gas then filtered with 0.22 um PES filters. Subsamples were stored at −20 °C for sulfate measurement, preserved with 0.5 M zinc acetate for sulfide analysis, or derived with monobromobimane (mBB) for sulfide, thiosulfate, and sulfite preservation [54]. For mBB derivatization directly following filtration, 50 μL of sample were thoroughly mixed with 50 μL of 60 mM mBB in acetonitrile and 75 μL buffer (50 mM HEPES; 5 mM EDTA; pH 8). The mixture reacted in the dark for 1 hr then was quenched with 200 μL of 65 mM methanesulfonic acid and frozen until analysis.

Sediment cores collected for microcosm incubations were sectioned in 1-3 cm horizons and 2 mL of sediment from each horizon was collected in a sterile cutoff syringe and frozen in a cryovial at -80 °C for molecular analysis. The remainder of each horizon was placed in heat-sealed mylar bags and flushed with N_2_ for ∼5 min to remove oxygen in the headspace. Sediments were stored at 4 °C until incubations were initiated in the lab (<60 days after collection).

### Microcosm setup and sampling

A representative sediment core (DR1171-LC63) was collected from ‘SW Clam Patch 1’, a methane seep encircled by vesicomyid clams and characterized by the presence of a yellow microbial mat. The sediment core was collected from the center of the seep site, where clams were absent.

Sediment sections within to below the zone of sulfate reduction (24-36 cm) were homogenized and suspended in 500 ml of filter-sterilized N_2_-sparged bottom water from the site in a 1:3 ratio. The slurry was split into two sections, one of which remained in natural seawater (NSW) collected from the site and a second which was washed three times with anoxic sterilized sulfate-free artificial seawater (SF-ASW) [55]. Microcosms were prepared on ice in 125 mL sterile serum bottles and capped with NaOH-washed blue butyl rubber stoppers. Serum bottles were flushed with N_2_ and filled with 60 mL of either the NSW slurry or the SF-ASW slurry. Prior to stoppering the bottles, 200 mg of elemental sulfur (Sigma-Aldrich, 100 mesh) was added to select microcosms, serving as the primary sulfur source. Following capping, microcosms were flushed for ∼5 min with either N_2_ or CH_4_ then over-pressurized (2 bar).

Freshly prepared, filter-sterilized, anoxic sulfur sources were then added to the appropriate microcosms to reach final concentration of either 30 mM sulfate or 10 mM thiosulfate. All microcosm treatments were conducted in duplicate. Microcosms were rotated in the dark at 4 °C and sampled after 14, 33, 62, 97, 154, 243, 434, and 650 days. During sampling, microcosms were kept on ice and shaken prior to sample collection to ensure homogenous slurry removal. Using an N_2_-flushed syringe, slurries were collected and immediately frozen for molecular analysis. Within an anaerobic chamber, slurries were collected by syringe and filtered with 0.22 um PES filters for analysis of dissolved sulfate, sulfide, thiosulfate, and sulfite. Filtered samples were preserved in the same manner as sediment core porewaters.

### Geochemical analysis

Sulfate was quantified by either ion chromatography (IC) or high-pressure liquid chromatography (HPLC). Sulfate was determined in porewater samples from sediment cores by IC using a Dionex ICS-2000 system (Dionex; CA, USA) as previously described [56]. Briefly, samples were diluted 1:50 with 18 MΩ water and anions were separated on a 2mm Dionex IonPac AS19 analytical column protected with a 2mm Dionex IonPac AG19 guard column (Thermo Fisher Scientific; MA, USA). A hydroxide gradient was supplied with a KOH generator cartridge and pumped (0.24 mL min^-1^) at 10 mM for 5 min, linearly increased to 48.5 mM at 27 min, and again linearly increased to 50 mM at 40 min with a Dionex AERS 500 (recycle mode, 30 mA) providing suppressed conductivity detection. Sulfate was quantified in incubations by HPLC (Thermo/Dionex UltiMate 3000) using a Metrohm Metrosep A supp 5 anion exchange column (150 mm × 4 mm) with 3.2 mM Na2CO3/1.0 mM NaHCO3 (0.7 mL min^-1^) eluent and absorbance measured at 215 nm [57]. To prevent oxidation of sulfur intermediates to sulfate, all dilutions for HPLC analysis were done the same day as measurements with N_2_ degassed solutions and sulfate measurements were compared in frozen and zinc acetate amended subsamples.

Sulfide preserved in zinc acetate was measured via a Cline method modified for use in a 96-well plate reader (Tecan Sunrise 4.2) [56]. Sulfide concentrations were confirmed, and thiosulfate and sulfite concentrations were quantified in mBB derived samples on an ACQUITY I-Class UPLC system (Waters, MA, USA) with an ACQUITY UPLC BEH C18 column (2.1 mm x 50 mm, 1.7 um, 130 A) held at 45 °C with 1 µL injection volume. The samples were eluted with A) 0.1% formic acid (LCMS grade), 99.9% MQ water and B) 100% ACN (LCMS grade). Gradient elution (0.4 mL min^-1^ throughout) began with 95% A, 5% B, decreased linearly following: 93% A at 1.75 min, 81% A at 1.8 min, 78% A at 3.5 min, and 5% A at 3.55 min. The gradient was maintained at 5% A until 3.95 min, then linearly increased to 95% A at 4 min, which was maintained until the run was completed at 4.5 min. Signal was quantified using an ACQUITY fluorescence detector (ex. 380 nm, em. 480 nm) and sulfur speciation confirmed using a Xevo G2-S TOF mass spectrometer (Waters; MA, USA) using previously optimized settings [54].

### Sulfur isotope analysis

Sulfur stable isotopes (^34^S/^32^S) were analyzed with an EA IsoLink combustion elemental analyzer system coupled to a Delta V Plus isotope ratio mass spectrometer (Thermo Scientific; Bremen, Germany) [58]. Sulfide samples preserved with zinc acetate were washed three times with deionized water, aliquoted appropriately, and dried in sulfur-free tins. Solid elemental sulfur and solutions of thiosulfate and sulfate were also aliquoted in sulfur-free tins for analysis. Calibration was achieved with SO_2_ reference gas peaks within each EA run, and by comparison to IAEA reference materials S1, S2, and S3; the analytical precision was better than ±0.3‰. No blank correction was required as SO_2_ blanks were <<1% of peak area. Sulfur isotope ratios, expressed as part-per-thousand (permil, ‰) deviations from the VCDT reference, were reported in the conventional delta notation:

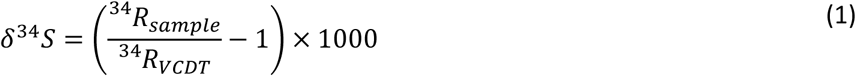

The isotopic fractionation was calculated as:

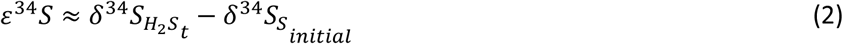

was determined at each timepoint (*t*) and *δ*^34^*S_S__initial_* was the isotopic ratio of the amended sulfur source.

### DNA extraction, amplicon sequencing, and analysis

DNA from slurry incubations and field samples was extracted with the DNeasy PowerSoil Pro Kit (Qiagen 12888; Hilden, Germany) according to the kit protocol, with an added bead beating step (45 s, 5.5 m/s; FastPrep FP120, Thermo Fisher Scientific; MA, USA). The V4-V5 region of the 16S rRNA gene was PCR amplified in duplicate using archaeal/bacterial primers 515f (5’- TCGTCGGCAGCGTCAGATGTGTATAAGAGACAGGTGYCAGCMGCCGCGGTAA-3’) and 926r (5’- GTCTCGTGGGCTCGGAGATGTGTATAAGAGACAGCCGYCAATTYMTTTRAGTTT-3’) with Q5 Hot Start High Fidelity 2x Master Mix (New England Biolabs, MA, USA). Following 30 annealing cycles at 54 °C, PCR duplicates were pooled and amplified with Illumina Nextera XT index 2 primer barcodes using Q5 Hot Start PCR mixture with 10 cycles at 66 °C. Purification was conducted with a MultiScreen Plate (Millipore-Sigma MSNU03010; MO, USA) with vacuum manifold and quantified with the QuantIT PicoGreen dsDNA Assay Kit (Thermo Scientific P11496; MA, USA) on a Touch Real-Time PCR Detection System (BioRad; CFX96). Equimolar amounts of barcoded samples were combined and purified (Qiagen; PCR Purification Kit 28104) and submitted to Laragen (Culver City, CA) for PE250 sequencing on Illumina’s MiSeq platform with addition of 15-20% PhiX control library.

Raw Illumina iTag reads were processed using DADA2 [59]. Sequences were trimmed (240f/200r) and merged with a 12 bp overlap. Chimeras were removed and taxonomy was assigned via IDTAXA [60] using SILVA 138.1 [61] amended with 1,197 in-house high-quality, methane seep-derived bacterial and archaeal clones (available upon request). Phylogeny for 16S rRNA genes of interest which were unclassified at the family or genus level was further investigated with ARB 6.0.6 [62], briefly, sequences were aligned to the Silva database and added to the existing tree by parsimony.

### Metagenomic and Metatranscriptomic sequencing and analysis

Samples from the final timepoint (650 days) were collected within an anaerobic chamber and immediately extracted using the Zymo Quick-DNA/RNA Microprep Plus Kit (Zymo Research Corporation, D7005). Metagenomic and metatranscriptomic libraries were prepared with NEBNext Ultra II DNA Library Prep Kit for Illumina (NEB, E7645) and the NEBNext Ultra II RNA Library Prep Kit for Illumina (NEB, E7770) respectively at the Caltech Genetics and Genomics Laboratory (Pasadena, CA, USA). Metagenomic libraries were sequenced by Novogene (HiSeq X Ten, PE150) resulting in an average of 31.9 million raw read pairs per sample (range: 26.8 – 40.1 million reads); trimming resulted in an average of 28.1 million read pairs per sample (range: 23.3 – 35.5 million reads). Transcriptomic libraries were sequenced by Novogene (NovoSeq 6000, PE150) resulting in an average of 93.5 million raw reads per sample (range: 72.3 – 118.2 million reads); trimming resulted in an average of 89.4 million read pairs (range: 111.7 – 69.9 million read pairs).

Metagenomic short reads were filtered for quality [63, 64] and samples from duplicate microcosm treatments were assembled using MEGAHIT (1.2.9) [65]. Gene calling was done with Prodigal (2.6.3) [66] and predicted genes were annotated using PFAM [67], COG [68], and KEGG [69] via Anvi’o wrapper scripts [70]. Transcriptomic data was indexed to deduplicated metagenomic gene calls from co-assembled treatments using KALLISTO [71], filtered by E-values < 10^-15^, and transcripts per million (TPM) for all transcripts were collated. Phylogenetic composition from transcriptomic data was determined with phyloFlash [72]. Genes of interest identified by KEGG Orthology were aligned to a curated database of genes derived from the non-redundant and taxonomically divergent database of bacteria and archaea in the Genome Taxonomy Database (GTDB r202) [73] using MUSCLE [74]. The databases were created with the use of either HMMs from MagicLamp [75] or custom-made for this purpose. The custom-made HMMs were made using either HMMer [76] from a filtered set of reference sequences available in an updated COG database [77] of NCBI or with a taxonomically divergent set of sequences from the Integrated Microbial Genomes (IMG) database [78]. All custom-made HMMs are available at https://github.com/ranjani-m/Sulfur_chemistry. Phylogenetic trees were inferred using IQ-TREE2 [79] with models selected automatically by Model Finder and branch support values were calculated with 1000 ultrafast bootstraps. Visualization of putative genes enabled identification of incorrectly called clades, which were removed from *phsA, soxY*, and *soxC* trees.

## Results and Discussion

### Monterey Methane Seeps are Active and Diverse

Within seep sites, high sulfide concentrations indicate active biogeochemical cycling (Fig. 1).

**Figure 1.**
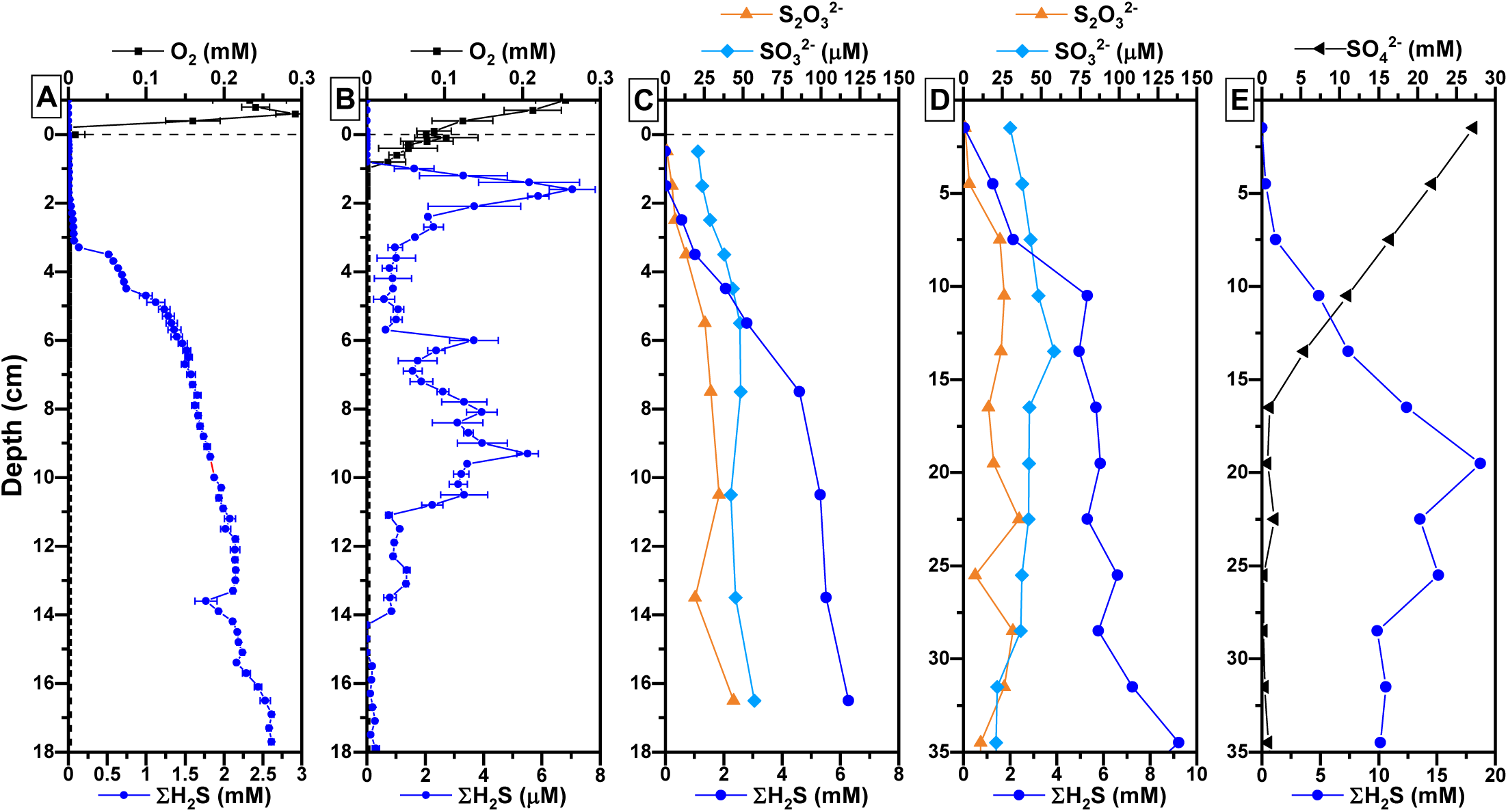
Depth profiles of dissolved oxygen, sulfide, sulfate, and sulfur redox intermediates in sediment cores. (A) Microelectrode depth profiles of oxygen and sulfide inside an active seep (DR1170-PC76) and (B) outside the seep (∼4 m away, DR1170-PC60). (C–D) Concentrations of sulfide, sulfite, and thiosulfate (measured by mBB) in active seep cores (DR1170-PC50, DR1170-LC67). (E) Sulfide (Cline assay) and sulfate (IC) from an active seep core (DR1030-LC46). Note the µM scale used for sulfide in B and longer profile depths obtained in D and E.

High-resolution depth profiles of one seep-site sediment core showed oxygen depletion at the sediment water interface (SWI) and sulfide concentrations up to 2.6 mM by 18 cm depth (Fig. 1A). Low sulfide accumulation (< 20 μM) within the top 2 cm can be attributed to sulfide-oxidizing microbial mats, as previously seen at mat-coved Monterey Canyon seeps [45] and other methane-rich environments [80]. In contrast, microelectrode profiles of sediment from the seep periphery showed oxygen penetration down to 8 mm and sulfide concentrations < 7.5 μM throughout the core (Fig. 1B). Lack of a visible microbial mat on the seafloor along with low sulfide concentrations indicate that outside of the seep area the rate of sulfate reduction is drastically diminished and the upward advection of methane is likely to be low [81, 82].

Extracted porewaters from active seep sites showed maximum sulfide concentrations of 6.3 mM, 9.2 mM, or even 19.5 mM, which corresponded with complete removal of sulfate by 25.5 cm (Fig. 1C,D,E). Despite the high sulfide concentrations and presence of sulfide-oxidizing microbial mats, sulfite and thiosulfate concentrations remained < 50 μM and did not increase at the SWI or appear to be dependent on sulfide concentrations (Fig. 1C,D).

Microbial community composition frequently varies with depth in active seeps, as documented in sediment core DR1171-LC63 used to seed microcosm incubations (Fig. 2) and observed in sediment cores from nearby seep sites (data not shown). Alpha diversity (Shannon index) remained high throughout the core, with a slight decline in zones of elevated ANME abundance. Both ANME-2c and ANME-1b, which commonly co-occur in methane seeps [81, 83], were abundant between 10.5-43.5 cm, whereas only ANME-1b persisted below 43.5 cm (Fig. S1A). *Desulfobacteraceae* group SEEP-SRB2 and to a lesser extent SEEP-SRB1, both known sulfate-reducing syntrophic partners of ANME [1, 84], were also observed throughout the core (Fig. S1B). Sulfur-oxidizing Campylobacterota, including *Sulfurovum* and *Sulfurimonas*, were concentrated in the upper horizons but remained detectable at depth, suggesting their activity extends into anoxic layers (Fig. S1C). However, of the bacterial lineages, members of the candidate phylum Caldatribacteriota JS1 were amongst the most abundant ASVs (Fig. 2). Members of JS1 are commonly observed in seep sediments [85, 86] and correlated well with ANME-2 in Nyegga cold seeps [83], although details around syntrophic relationships remain uncertain. Such correlations paired with the high diversity found in these sediments may suggest the potential for alternative partnerships (e.g. [87]) and reduction of electron acceptors other than sulfate in seep sites [88].

**Figure 2.**
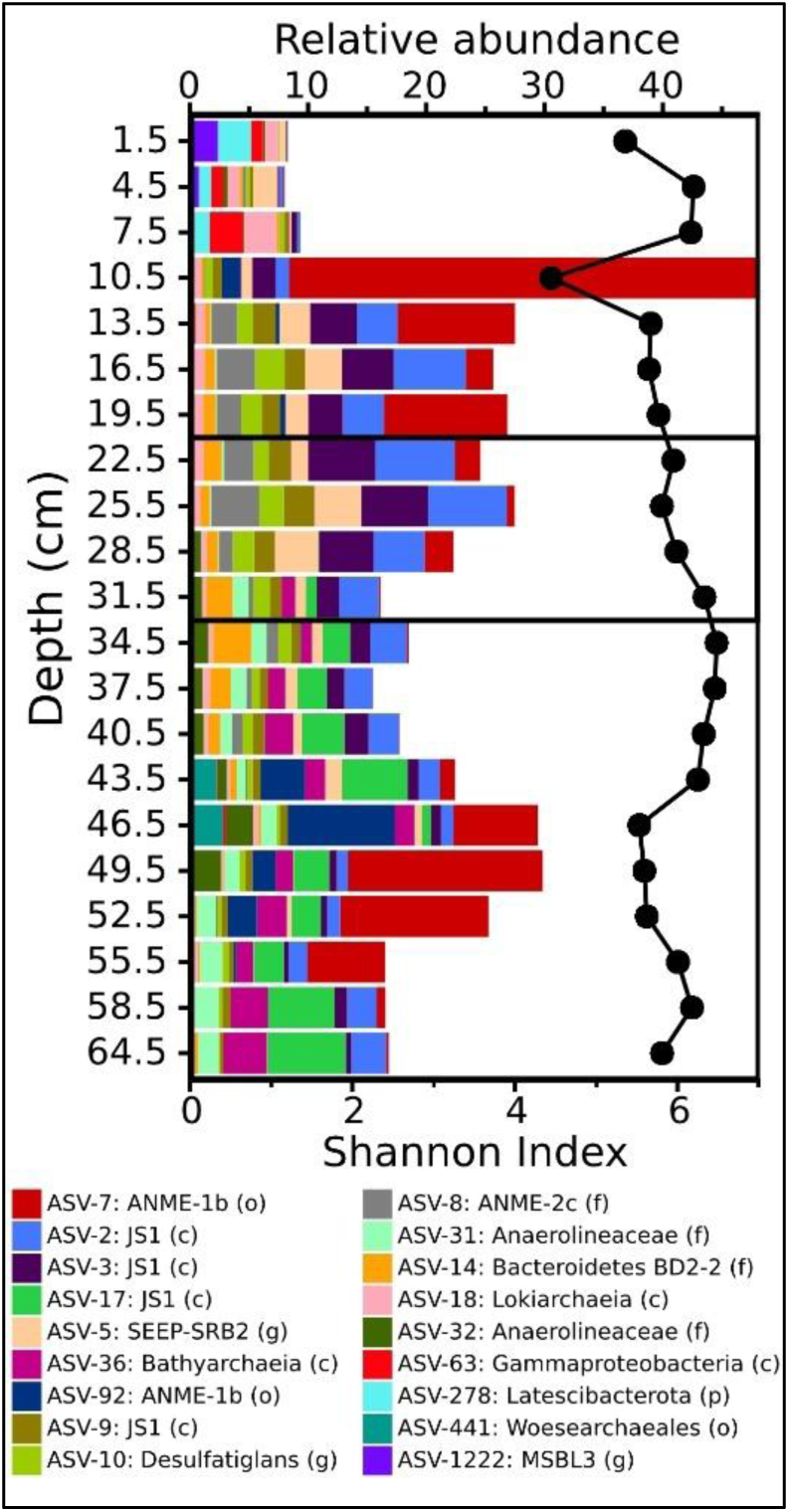
Microbial community composition with depth in seep sediment core DR1171-LC63. Shown are relative abundances of 16S rRNA ASVs >3%, the Shannon diversity index, and horizons used to seed microcosm incubations (black box).

### Sulfur source influences bacterial, but not archaeal community composition

Anoxic microcosms were established with sediment from 24-36 cm (black box, Fig. 2) and amended with one sulfur source (elemental sulfur, thiosulfate, or sulfate), with or without methane. Without the addition of organic electron donors, turnover rates within the original sediments are expected to be low [26], necessitating long incubation times. Still, over the 650-day incubation period, Shannon diversity remained high in all treatments, consistent with the original sediment core.

Inspection of the Desulfobacterota revealed persistence of specific SEEP-SRB2 lineages (ASV-5, ASV-35) from the original sediment core (Fig. 3A, S2). Sulfur treatment did not impact SEEP-SRB abundances except for ASV-882, which increased modestly in thiosulfate amended microcosms following day 153 (Fig. S3). Members of the *Desulfocapsaceae*, some of which are known sulfur disproportionators [24], also showed preference for thiosulfate. Several *Desulfacapsa* lineages (ASV-112, ASV-293, ASV-357) increased transiently, while ASV-28 (98.1% similarity to SEEP-SRB4) dominated thiosulfate treatments by day 242. Other Desulfobacterota also varied over time: after day 97, an uncultured Desulfobacterales (ASV-57) increased in thiosulfate and element sulfur treatments, while a *Desulfonatronobacter* (ASV-154) rose in sulfate treatments after day 153, especially without methane. In contrast, sulfur or methane amendments had no significant effect on the composition and proportion of archaeal Halobacterota, with all incubations retaining a relatively high abundance of the ANME-2c (ASV-8) and ANME-1b (ASV-7) lineages documented in the original sediment (Fig. 3B, S4).

**Figure 3.**
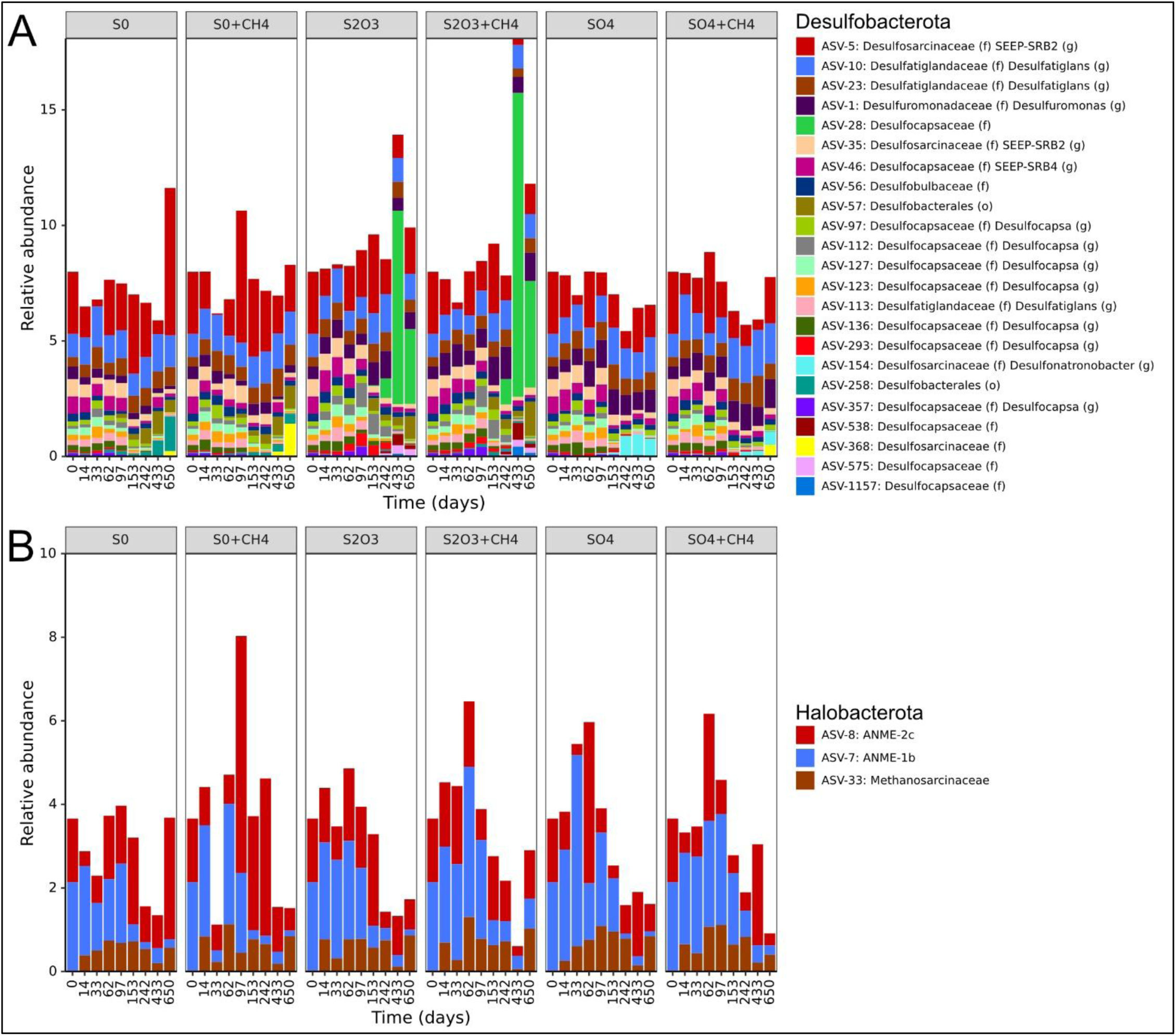
Dynamics of key microbial lineages in microcosms over 650 days. (A) Desulfobacterota ASVs (>0.4% relative abundance) and (B) Halobacterota ASVs (>1.0% relative abundance). Note that ASV designations match Figure 2.

### Geochemical Evidence for Complex Sulfur Cycling

Despite similarities in the microbial community composition between incubations, clear differences in sulfide production were observed (Fig. 4). Interestingly, initial rates of sulfide production were highest in microcosms with solid elemental sulfur, producing 2.6 ± 0.2 mM sulfide after 650 days. Thiosulfate amendments produced 2.3 ± 0.2 mM sulfide by day 650, while sulfate and natural seawater amendments produced 0.26 ± 0.06 mM and 0.16 ± 0.01 mM sulfide respectively by the ultimate timepoint. Sulfide concentration is dependent on electron acceptor oxidation state such that more reduced sulfur sources should produce more sulfide per mole carbon oxidized. For example, with acetate as electron donor, elemental sulfur reduction yields 2 mols sulfide per carbon, while thiosulfate yields 1 mol and sulfate only 0.5 mol sulfide per carbon (Eq. 1-3).

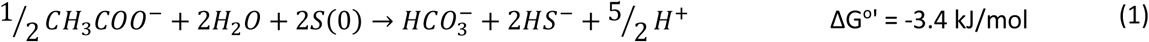

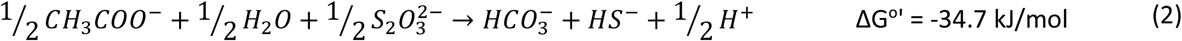

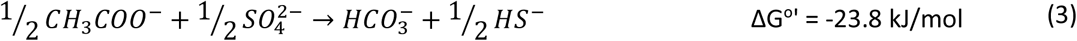

**Figure 4.**
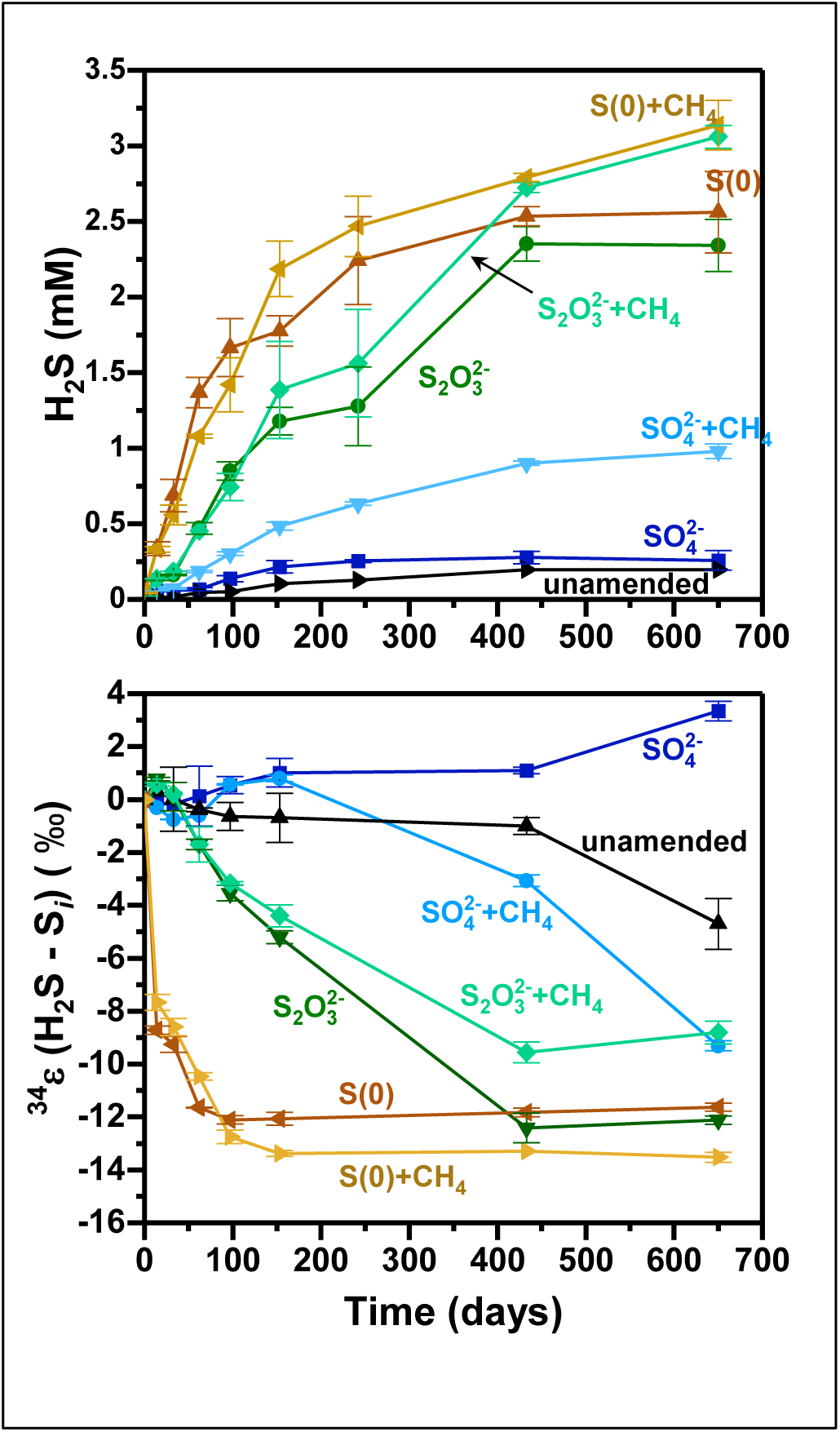
Geochemical and isotopic patterns in anoxic microcosms over time. (A) Sulfide concentrations in treatments amended with different sulfur sources and methane conditions. (B) Sulfide isotopic fractionation (ε³⁴S), calculated relative to the δ³⁴S of the initial sulfur source.

However, oxidation state alone cannot explain the variation in sulfide concentrations observed as greater than 4x sulfide was produced with elemental sulfur compared to the sulfate condition and > 2x sulfide was measured in thiosulfate treatments relative to sulfate. These discrepancies suggest that additional pathways contribute to sulfide accumulation, highlighting the complexity of sulfur cycling in deep-sea sediments. One such pathway is the disproportionation, or inorganic fermentation, of sulfur redox intermediates, which can simultaneously generate sulfide and sulfate without requiring an organic carbon source. While the disproportionation of thiosulfate is thermodynamically favorable (Eq. 4), elemental sulfur disproportionation (Eq. 5) is endergonic at sulfide concentrations > 1 mM [24] and cannot sustain bacterial growth unless sulfide is removed [89].

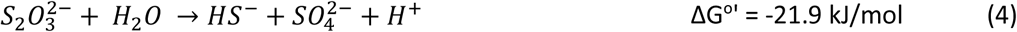

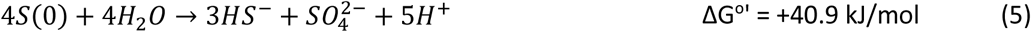

High sulfide concentrations and the lack of sulfide scavengers (e.g. ferrihydrite) argue against elemental sulfur disproportionation. Yet, rapid reaction between elemental sulfur and sulfide could allow for polysulfide disproportionation, as proposed in some ANME-2–SRB consortia [90].

Given the influence of sulfur source on sulfide accumulation, the effects of methane addition were also examined. In all cases, methane amendment increased sulfide yields, with elemental sulfur and thiosulfate treatments reaching similar final concentrations of 3.2 ± 0.2 and 3.1 ± 0.1 mM respectively, while the sulfate treatment only produced 0.98 ± 0.05 mM sulfide after 650 days (Fig. 4). In marine sediments, AOM is classically coupled to sulfate reduction, however, thiosulfate- or sulfite-driven AOM is predicted to be more energetically favorable (Eq. 6-8). Although elemental sulfur-driven AOM is endergonic under standard conditions (Eq. 9), polysulfide-driven AOM may be more favorable and has been proposed under sulfate-limiting conditions [40].

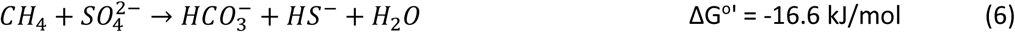

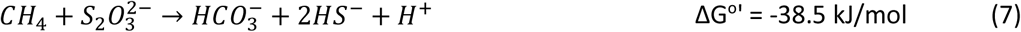

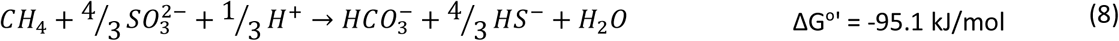

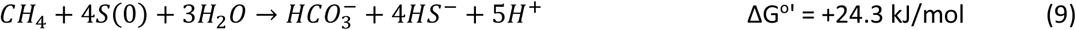

While methane addition consistently increased sulfide production, its effect on other sulfur compounds varied. Sulfite was only detected in thiosulfate-amended microcosms and increased modestly over time with no impact due to methane (Fig. S5). Similarly, thiosulfate consumption showed little change despite higher sulfide concentrations with methane addition. In contrast, sulfate concentrations were lower in sulfate-plus-methane and natural seawater treatments than in sulfate-only incubations, consistent with the higher sulfide concentrations in those treatments. Both thiosulfate and sulfate were below detection in microcosms not initially amended with those sulfur sources.

Notably, the absence of sulfate in elemental sulfur- and thiosulfate-amended microcosms may challenge disproportionation or point towards rapid reduction following sulfate production. Overall, these geochemical patterns highlight the need for interdisciplinary methods, such as sulfur isotope analysis and transcriptomics, to fully resolve the complex sulfur cycling in these sediments.

### Sulfur and methane treatments differentially influence sulfur isotopic fractionation

In addition to impacting sulfide concentrations, sulfur and methane treatment had a clear effect on the isotopic fractionation (ε^34^S) of sulfide (Fig. 4). In both treatments with sulfate as the sole sulfur source and in the natural seawater incubation where sulfate was abundant, ε^34^S values remained small (±1 ‰) through day 433. ^34^S-depletion of sulfide is common during organoclastic sulfate reduction, however bacterial species and environmental conditions can cause ε^34^S to span from the theoretical limits of depletion (∼75 ‰) to slight enrichment (∼3 ‰) (e.g. [91]). By the final timepoint, ε^34^S was +3.3 ± 0.4 ‰ in the ASW-sulfate treatment, while a ε^34^S of -4.7 ± 0.9 ‰ was observed in the natural seawater treatment. While fractionation was minimal in sulfate-amended microcosms, ε³⁴S exhibited a larger and earlier decrease in the elemental sulfur– and thiosulfate–amended microcosms. In elemental sulfur microcosms without methane, ε³⁴S dropped -9.3 ± 0.3 ‰ in the first 33 days and reached −11.6 ± 0.2 ‰ at the final timepoint, consistent with values reported for both reduction and disproportionation of elemental sulfur. Isotopic investigation of isolated elemental sulfur reduction is limited, however, cultured *Dethiosulfovibrio* species produce ^34^S-depleted sulfide (-1.3 to -5.2 ‰ [92]). Sulfide from elemental sulfur disproportionation, which should initially proceed through a similar mechanism as elemental sulfur reduction, ranged from −6.2 to -8.0 ‰ in sediment enrichments, -3.7 to -5.9 ‰ in cultures of *Desulfocapsa* species, and −15.5 ‰ in *Desulfobulbus propionicus* cultures [93, 94].

Thiosulfate-amended microcosms exhibited a comparable final ε³⁴S, but with a steady decline to −12.1 ± 0.2 ‰ by day 433, matching prior reports of thiosulfate reduction and disproportionation (-7 to −12 ‰ [21, 22, 95].

While sulfide production increased in all treatments with methane addition, isotopic responses diverged. In sulfate-plus-methane microcosms, ε³⁴S decreased to -9.3 ± 0.2 ‰ by day 650, lower than in sulfate-only controls, but within the upper bounds of fractionation in other seep sediments [96]. Data from AOM cultures is limited, but previous reports show sulfate-driven AOM produces fractionations that overlap with organoclastic sulfate reduction [97]. Although complete sulfate removal in the SMTZ may produce ^34^S-enrichment of sulfide minerals [32], this cannot explain the relatively mild fractionation observed in these microcosms as sulfate remains in excess (Fig. S5) and Rayleigh distillation effects would be minimal. The supplied sulfur compounds also remained abundant in elemental sulfur- and thiosulfate-amended microcosms. Like sulfate-amended microcosms, the increased sulfide accumulation in elemental sulfur-plus-methane treatments was accompanied by slightly greater fractionation at the final timepoint (-13.5 ± 0.2 ‰) relative to the paired microcosm without methane. In contrast, thiosulfate-plus-methane microcosms produced an additional 0.75 mM sulfide but had a positive isotopic shift relative to methane-free microcosms, reaching only -8.8 ± 0.4 ‰.

Together, these results highlight the strong influence of both sulfur source and electron donor on isotopic systematics in deep-sea sediments. The relatively small fractionations across treatments suggest complex sulfur cycling, potentially including disproportionation or oxidation of sulfur redox intermediates. The contrasting ε³⁴S responses to methane in thiosulfate versus sulfate and elemental sulfur treatments further underscore process complexity. While extreme fractionations in marine sediments have traditionally been attributed to oxidation and disproportionation [98, 99], cryptic sulfur cycling has more recently been recognized as a mechanism that can attenuate isotopic signals in sediments harboring diverse microbial communities [100]. The specific biogeochemical pathways contributing to sulfur cycling in these sediments are evaluated below based on an integration of the geochemical, isotopic, and transcriptomic results.

### Seep-associated Desulfobacterota and Campylobacterota demonstrate disproportionation potential

Disproportionation of thiosulfate and elemental sulfur is initially catalyzed by a molybdopterin-dependent enzyme, annotated as either *phs* (signifying *p*roduction of *h*ydrogen *s*ulfide) related to thiosulfate reduction, or *psr* (signifying *p*oly*s*ulfide *r*eduction) more commonly referred to during elemental sulfur reduction. Both catalytic subunits *phsA* and *psrA* share sequence similarities [101] and are denoted by the same KEGG Orthology (K08352), thus we employ ‘*phs’* to refer to both. Bulk *phsA* transcripts were higher in thiosulfate treatments regardless of methane addition (Fig. 5A) and phylogenetic analysis showed increased expression by the *Desulfocapsaceae* and *Desulfobulbaceae* in thiosulfate amended experiments (Fig. 6A, S6). Members of both families have been shown to grow during either thiosulfate or elemental sulfur disproportionation [24]. In *Desulfocapsa sulfexigens*, sulfide production during thiosulfate disproportionation was attributed to thiosulfate reductase, whereas the mechanism during elemental sulfur disproportionation could not be resolved [102]. Similarly, in seep microcosms, *phsA* upregulation and increased *Desulfocapsaceae* 16S rRNA abundances occurred with thiosulfate amendment but not elemental sulfur, suggesting that elemental sulfur reduction and/or disproportionation involves alternative enzymes or taxa.

**Figure 5.**
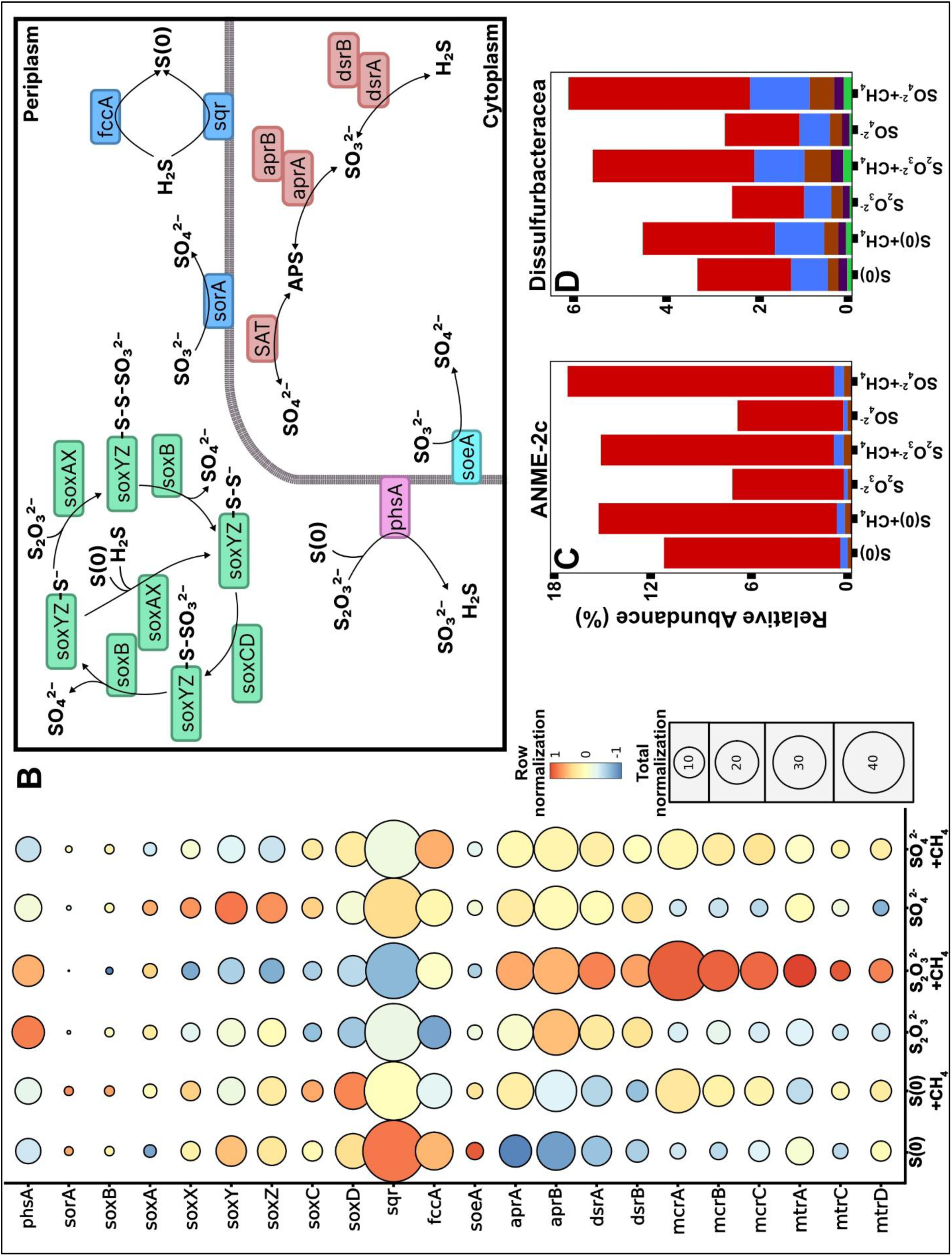
Transcriptomic evidence of sulfur and methane cycling. (A) Bulk transcript abundances for sulfur and methane metabolism genes, normalized by treatment (color) and overall transcript abundance (size). (B) Schematic of periplasmic and cytoplasmic sulfur cycling pathways. (C) Relative SSU rRNA transcription of ANME-2c and (D) SEEP-SRB2.

**Figure 6.**
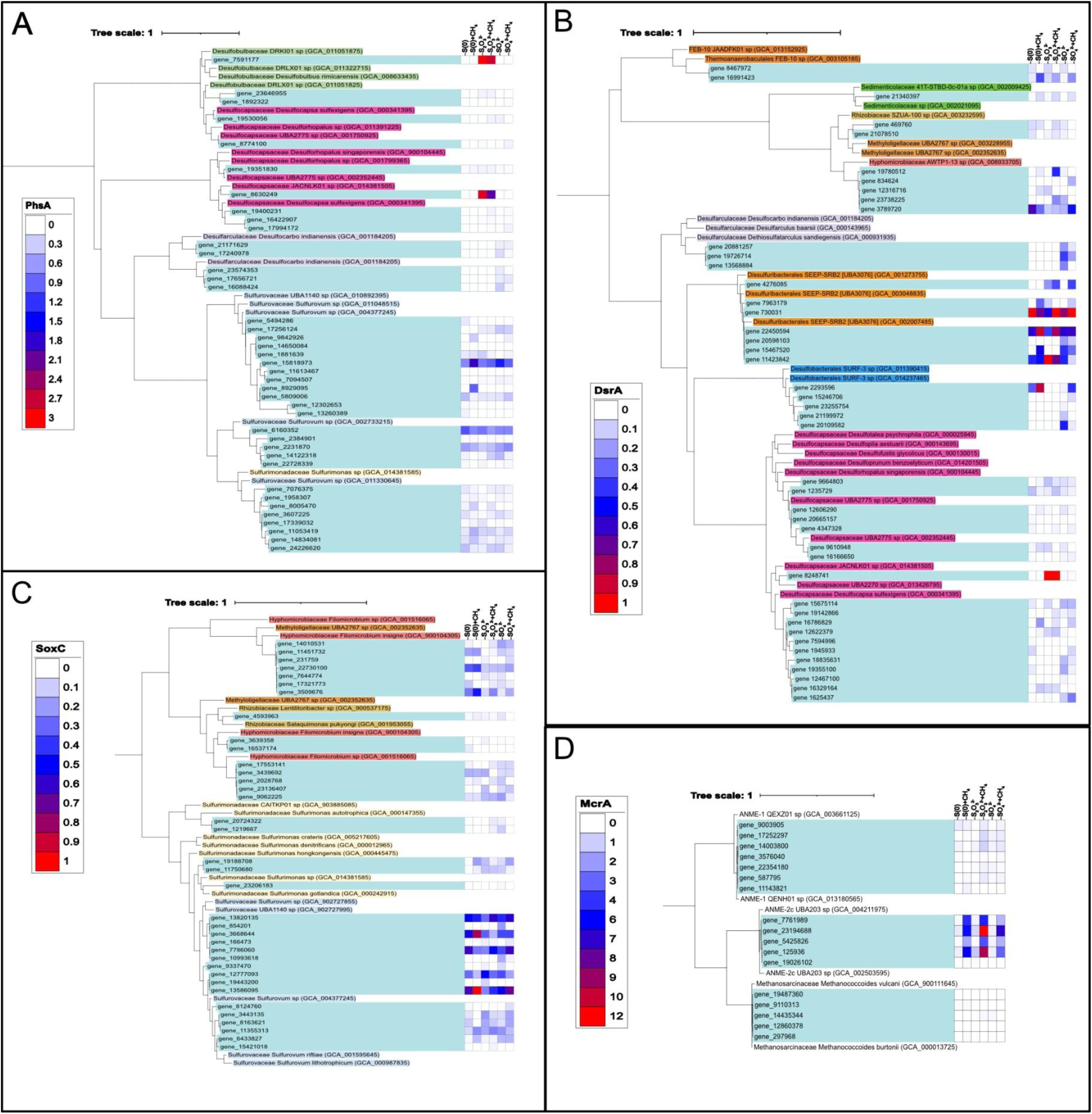
Phylogenetic placement of key functional genes expressed in seep microcosms. Shown are trees for (A) *phsA*, (B) *dsrA*, (C) *soxC*, and (D) *mcrA*.

Sulfur-source dependent expression of phsA was also observed in the *Desulfarculaceae* (Fig. 6A). Despite being a known sulfate, sulfite, and thiosulfate reducer [103], *Desulfarculaceae* only expressed *phsA* in sulfate treatments. In contrast, *phsA* transcripts associated with members of the Campylobacterota, *Sulfurovaceae* and *Sulfurimonadaceae,* were upregulated in all treatments (Fig. 6A).

These lineages have been proposed to disproportionate sulfur redox intermediates [104], and their prevalence in deep-sea vent environments has been linked to elemental sulfur reduction via *phs* coupled to H_2_ oxidation [105–107].

Importantly, *phs* can support thiosulfate or elemental sulfur reduction independent of disproportionation [20, 101, 108, 109]. During thiosulfate cleavage by *phs,* sulfide and sulfite are produced within the periplasm [110, 111] and the extracellular release of sulfite has been confirmed in both thiosulfate-reducing and disproportionating cultures [102, 108]. However free sulfite reacts abiotically with sulfide to reform thiosulfate [112] and complete removal of sulfite has been observed in disproportionating cultures with sulfide addition [110]. Thus, the gradual sulfite accumulation in thiosulfate-amended microcosms despite much higher sulfide concentrations likely reflects either enhanced thiosulfate cleavage or reduced microbial sulfite removal. Together, *phs* upregulation, family-specific expression patterns, and sulfite accumulation may indicate a potential for thiosulfate disproportionation, however, these observations only account for the reductive half of this process.

### Genes involved in periplasmic sulfur oxidation are transcribed in all treatments

Complete disproportionation requires an oxidative step yielding sulfate. For thiosulfate, this likely proceeds through sulfite, while in elemental sulfur disproportionation the oxidized intermediate remains unresolved. Sulfite transformation within the periplasm would minimize transport demands and limit the redox stress associated with cytoplasmic accumulation [23]. Induction of the periplasmic sulfite oxidase *sorA* (Fig. 5B) is reported with thiosulfate in prominent sulfite oxidizers *Starkeya novella* and *Sinorhizobium meliloti* [113] and *sorA* activity, as determined by a ferricyanide-dependent assay, was demonstrated in disproportionating *D. sulfexigens* cultures [102]. However, *sorA* is absent in other Desulfobacterota and later studies failed to identify *sorA* within the *D. sulfexigens* genome [114]. By contrast, *sorA* has been detected in *Sulfurovum* and *Sulfurimonas* isolates [115] and may facilitate disproportionation by these lineages [104]. Still, *sorA* transcription was low in all microcosms and especially minimal with thiosulfate amendments (Fig. 5A), suggesting that this is not a dominant mechanism for sulfite oxidation in these seep sediments.

Sulfur oxidation within the periplasm may also proceed via the sulfide quinone oxidoreductase (*sqr*) or flavocytochrome c (*fcc*). Despite the relative upregulation of these genes in all microcosms (Fig. 5A), their accurate annotation is challenging as multiple closely related homologs are known to perform a wide range of functions (e.g. glutathione reduction, lipoamide dehydrogenation, or NADH reduction). As such, it is possible that some of the *sqr* and *fcc* kegg-annotated homologs detected in our seep microcosms are performing a different type of chemistry altogether. Furthermore, sulfur oxidation by *sqr* and *fcc* is primarily observed in phototrophs or chemotrophs with the main function of sulfide detoxification, resulting in elemental sulfur/polysulfide production rather than sulfate (Fig. 5B) [116]. Thus, *sqr* and *fcc* are unlikely to generate sulfate during disproportionation by *Desulfocapsaceae* or *Desulfobulbaceae*.

Finally, periplasmic oxidation of elemental sulfur, sulfide, and thiosulfate is commonly observed by the *sox* pathway [115, 117]. Although sulfite oxidation has only been demonstrated with purified *sox* enzymes [118, 119], the rapid abiotic reaction of sulfite with sulfide to form thiosulfate could support indirect *sox*-dependent oxidation. Within the dark, anoxic microcosms, transcripts for the complete *sox* pathway were detected (*soxCDYZXAB*), with the highest transcription observed in *soxY* and *soxZ*, likely due to their key sulfur-binding role (Fig. 5A). Both *soxY* and *soxZ* were relatively upregulated in thiosulfate and sulfate treatments without methane, whereas *soxC/D* decreased in thiosulfate amendments, consistent with their requirement for sulfate production during sulfide and elemental sulfur oxidation, but not during thiosulfate oxidation [117].

Phylogenies of *soxC* and *soxY* revealed broad transcription of these genes affiliated with members of the *Methyloigellaceae, Rhizobiaceae, Hyphomicrobiaceae*, and *Campylobacterota*, unaffected by sulfur source (Fig. 6C, S7, S8). No *sox* transcripts related to the Desulfobacterota were detected, confirming the well-known exclusion of *sox* and *dsr* genes [33, 120, 121]. While the abundance of *sox* transcripts affiliated with the *Sulfurovaceae* and *Sulfurimonadaceae* in these seep microcosms was unexpected, unlike many obligately phototrophic or (micro)aerophilic sulfur oxidizers, Campylobacterota are extremely adaptable. They have been observed in diverse sulfidic environments [122–124], and can anaerobically oxidize reduced sulfur species in conjunction with a variety of terminal electron acceptors including organic carbon, nitrate, nitrite, elemental sulfur, polysulfides, or thiosulfate [115, 125, 126]. Transcription of both *phs* and *sox* genes by the *Sulfurovaceae* and *Sulfurimonadaceae* may be indicative of separate reduction and oxidation processes, or potentially these two combined pathways may support disproportionation by these lineages in seep sediments. Overall, the minimal influence of sulfur treatment on transcription highlights the metabolic flexibility of these taxa and suggests that such microbial pathways play a larger role than previously expected in anoxic marine sediment.

### Disproportionation by Desulfocapsaceae is challenged by evidence of cytoplasmic sulfite reduction

In addition to periplasmic sulfur oxidation, the potential for cytoplasmic sulfate generation was evaluated. Transcripts of the sulfite-oxidizing gene *soeA* were detected in all treatments and slightly upregulated with elemental sulfur (Fig. 5A), however this pathway has not been identified in Desulfobacterota, and thus could only support disproportionation by Campylobacterota [127]. Alternatively, sulfite oxidation via the reverse functions of adenylyl-sulfate reductase (*Apr*) and ATP sulfurylase (*sat*) have been proposed in Desulfobacterota during disproportionation, despite the minimal energy yield [110]. *AprA* and *aprB* transcripts were consistently more abundant than *phsA and soeA* (Fig. 5A), and *aprA* phylogeny revealed strong upregulation in thiosulfate treatments from a lineage closely related to *D. sulfexigens* (Fig. S9). While *D.sulfexigens* cannot reduce sulfate despite encoding *aprA* [114], many other disproportionating members of the *Desulfocapsaceae* can, using *aprA* in the forward, reductive direction to reduce sulfate [128]. Given the divergence of sulfate-reduction capabilities within this family, exclusive reverse *Apr* function cannot be established based on transcripts alone. Furthermore, in thiosulfate-amended microcosms, *dsrA* and *dsrB* transcripts associated with a Desulfocapsaceae member closely related to *D. sulfexigens* were also strongly expressed (Fig. 6B, S10-S11). While these results could in principle suggest that one Desulfocapasaceae lineage was simultaneously oxidizing and reducing sulfite, upregulation of both *apr* and *dsr* could also reflect conventional sulfate reduction, with sulfate produced by other *sox*-containing community members. Further study of the seep-associated Desulfocapsaceae identified within these microcosms is required to explicitly determine their sulfate-reducing vs sulfite-oxidizing capacity.

### Abundant species level diversity contributes to sulfate reduction in all microcosms

Along with the specific Desulfocapsaceae-related transcripts discussed above, transcripts of the canonical sulfate reduction genes (*aprA, aprB, dsrA, dsrB*) were abundant in all microcosms (Fig. 5A), with a higher number of unique transcripts relative to *phsA* (Table. S2). Together with the 16S rRNA data (Fig. 3A), this indicates sustained diversity of putative sulfate reducers across all incubations. In elemental sulfur- and thiosulfate-amended microcosms, this diverse sulfate-reducing community likely contributes to the rapid consumption of any sulfate produced during sulfur oxidation or disproportionation, leading to sulfate concentrations below detection (Fig. S5).

Treatment-dependent expression of *aprA, dsrA*, and *dsrB* was also observed outside the *Desulfocapsaceae*. Transcripts associated with the *Desulfarculaceae* were upregulated in sulfate treatments, while those related to the uncultured Desulfobacterales *SURF-3* group were specific to both sulfate and elemental sulfur treatments (Fig. S9, 6B, S11). Although little is known about the *SURF-3* (renamed *Desulfosalsimonadaceae* in gtdb R214), they are related to the *Desulfonatronobacter* [129], which interestingly had higher 16S rRNA relative abundance in the sulfate microcosms (Fig. 3A).

Overall, the clade-level response to methane addition was minimal, although some taxon-specific transcripts were upregulated in the presence of methane based on *dsrA* phylogeny, including representatives from the Thermoanaerobaculales FEB-10 (gene_16991432), *Sedimenticolaceae* (gene_21340397), and the putative AOM syntrophic partner, SEEP-SRB2 (Dissulfuribacterales UBA3076, gene_22450594) (Fig. 6B). Increased transcription of all three sulfate-reduction genes (*aprA, dsrA, dsrB*) was associated with the SEEP-SRB2 clade, however, individual SEEP-SRB2 lineages differed between sulfur treatments. For example, in the *dsrA* phylogeny, one SEEP-SRB2 lineage (gene_11423842) was upregulated with thiosulfate while another (gene_20598103) was only transcribed in sulfate-amended microcosms. This transcript heterogeneity within the SEEP-SRB2 highlights broad species-level diversity in SMTZ sediment and suggests that different species from this clade may exploit different sulfur compounds or partner with different ANME lineages, as recently demonstrated for Seep-SRB1a [1].

### AOM is stimulated by thiosulfate and elemental sulfur amendments

With methane addition, sulfide concentrations increased similarly (580 −720 μM) across elemental sulfur, thiosulfate, and sulfate microcosms (Fig. 4A). Previous reports also demonstrated increased sulfide production rates with the addition of thiosulfate and methane to marine sediments, however minimal impact was observed with elemental sulfur-plus-methane additions [41, 43, 44]. Such variation likely reflect heterogeneity in seep microbial communities. Whereas earlier works relied on sediment stored for extended periods prior to investigation, the current study used fresh sediment from an active seep as the inoculum, where microbes may be more acclimated to fluctuating sulfur compounds and elemental sulfur accumulation [37, 38]. In contrast to prior enrichments showing reduced ANME and SEEP-SRB abundances with thiosulfate and elemental sulfur addition [41, 42], no such decrease was observed here over 650 days (Fig. 3). Instead, small subunit (SSU) rRNA transcription revealed increased activity of ANME-2c under methane addition across all sulfur sources (Fig. 5C).

ANME-1b activity was lower overall but also enhanced in thiosulfate and sulfate treatments (Fig. S12). For all sulfur sources, transcripts associated with AOM increased with methane addition (Fig. 5A), and *mcrA* phylogeny indicated ANME-2c as the primary contributor (Fig. 6D). SEEP-SRB2 showed parallel SSU rRNA expression patterns (Fig. 5D), consistent with its syntrophic role in AOM. Together, these results support the persistence of ANME-2c–SEEP-SRB2 syntrophy mediating AOM regardless of sulfur treatment. Interestingly, the increased ANME-2c SSU rRNA transcription in elemental sulfur microcosms lacking methane may point toward novel mechanisms of syntropy based on sulfur source. More broadly, the divergence between transcriptional activity and 16S rRNA gene persistence demonstrates the value of metatranscriptomic data for assessing microbial processes in these low-energy, deep-sea environments where microbes may be capable of both long-term dormancy and rapid adaptation depending on environmental conditions.

### Contextualizing sulfur isotopic fractionation with transcriptomic data

In light of transcriptomic trends, sulfide isotopic fractionation can help clarify cryptic process in these deep-sea sediments. Consistent transcription of the *sox* pathway indicates active sulfur oxidation regardless of treatment, however, the predominant oxidizable substrates varied. In sulfate-supplied microcosms, the absence of detectable sulfur redox intermediates implied that sulfide was the primary oxidizable substrate, enriching residual sulfide in ³⁴S through production of ³⁴S-depleted sulfate, as shown in *Sulfurimonas denitrificans* which expresses an isotropic fractionation of −2.4 to -3.6‰ [130].

Relative upregulation of *soxAXYZ* transcripts in the sulfate treatment without methane may indicate a larger proportion of sulfide oxidation relative to other microcosms, and indeed following day 153 sulfide concentrations stabilized, suggesting balanced sulfide production and oxidation. This stagnation may also reflect shifts in electron donor availability. Although slow sulfide production and electron donor limitations generally produce increased fractionation [131], electron donor identity is also important [132, 133]. Furthermore, slow sulfate-reduction rates do not necessarily imply reliance on low-quality electron donors, but may also arise from a slow supply of a high-quality donor such as H₂, which can support strongly irreversible sulfate reduction even when net rates appear low. Thus, organic carbon limitations may have increased the contribution of H_2_-driven sulfate reduction, which would not result in the same large fractionations expected with organic substrates at such low rates [134, 135]. Indeed, the divergence in ε^34^S between ASW-sulfate and natural seawater treatments at the final timepoint may reflect differences in electron donors as the natural seawater microcosms likely contained additional organic substrates that were inadvertently removed during the washing procedure invoked in ASW microcosms to eliminate dissolved sulfur species. In the ASW-sulfate treatment, sulfate reduction would have depended on electron donors retained during the washes or generated during incubation (e.g. H_2_). Transcriptomic data was not collected from natural seawater microcosms, however, in ASW-sulfate microcosms without methane, upregulation of hydrogenase genes supports H_2_ use (Fig. S13). Combined, the increased proportions of sulfide oxidation and H_2_-driven sulfate reduction contribute to the slight d^34^S_sulfide_ enrichment by the final timepoint in ASW-sulfate-amended microcosms.

With the addition of methane to ASW-sulfate treatments, *sox* transcription decreased, indicating reduced oxidation when electron donors were abundant. Instead, active AOM, as demonstrated by upregulation of *mcr* and *mtr* genes, reinforced the importance of electron-donor supply on the expressed isotopic fractionation associated with sulfate reduction in seep environments by altering the relative amounts of sulfide oxidation and production, ultimately shifting ε³⁴S to more negative values. Even so, fractionation remained modest compared to other marine sediments, suggesting that the sequential oxidation-disproportionation cycles traditionally associated with large ε³⁴S [98, 99] were not predominant within the timeframe of this study, despite clear transcriptomic evidence for sulfide oxidation. Elemental sulfur and thiosulfate microcosms can better isolate cryptic processes, but it is clear that more studies are needed to differentiate modern processes from the background isotopic patterns that accumulate over longer timescales in these environments.

In elemental sulfur microcosms, sulfide is not the most abundant reduced sulfur species, and instead the plentiful elemental sulfur likely served as the main oxidizable substrate for *sox*. Unlike sulfide oxidation, this would not result in ^34^S-enrichment of the sulfide pool. Still, oxidation of both elemental sulfur and sulfide are dependent on *soxC/D* (Fig. 5B) and likely produce similarly ^34^S-depleted sulfate (-2.4 to -3.6‰), whereas elemental sulfur disproportionation produces strongly ^34^S-enriched sulfate (+10.9 to +30.9 ‰ [93, 94]. The absence of sulfate accumulation (Fig. S5) and strong *apr/dsr* transcription indicate complete reduction of transient sulfate, producing sulfide with equivalent isotopic signals. While the complete reduction of sulfate produced via *sox* would only result in slightly ^34^S-depleted sulfide, reduction of the sulfate generated from disproportionation would produce significantly ^34^S-enriched sulfide. Accounting for the 3:1 ratio of sulfide to sulfate initially produced during disproportionation (Eq. 9), if disproportionation dominated, the resulting sulfide should have a more positive ε^34^S than the −12 ‰ observed over several timepoints in microcosms both with and without methane. Thus, disproportionation of elemental sulfur alone cannot explain the isotopic signals. This is even clearer with methane addition as, similar to sulfate-amended microcosms, ε^34^S was more negative than in microcosms without methane.

With thiosulfate amendment, sulfide was again not the dominant sulfur species available for oxidation via *sox*. However, unlike sulfide and elemental sulfur oxidation, initial sulfate production from thiosulfate oxidation does not require *soxC/D* (Fig. 5B), and indeed, these genes were not as upregulated (Fig. 5A). Although thiosulfate oxidation by photosynthetic microbes may produce little to no fractionation [136], chemolithoautotrophic oxidation has an inverse isotope effect and produces ^34^S-enriched sulfate (+2.9 to +5.8) [137, 138] due to the isotopic preference of sulfonate S in thiosulfate. Similarly, this internal isotopic preference contributes to sulfate enrichment (+12 ‰) during disproportionation [22]. Thus, the sulfate isotopic signals generated by either oxidation or disproportionation of thiosulfate are not as distinct as with elemental sulfur. Sulfate was again below detection and sulfate reduction genes were actively transcribed, indicating removal of transiently produced sulfate. However, given the similar isotopic signals of sulfate generated from *sox* and disproportionation mechanisms, any sulfide produced from reduction of this sulfate would be indistinguishable, making differentiation of the two pathways based on isotopic signals more challenging. In contrast to elemental sulfur-amended microcosms, the addition of methane does not provide much insight into the possibility of disproportionation, as ε^34^S was less depleted than in microcosms without methane.

### Potential Mechanisms for Anaerobic Oxidation of Methane

AOM in thiosulfate- and elemental sulfur-amended microcosms could be attributed to three distinct mechanisms. First, AOM may be coupled to the direct reduction of thiosulfate or elemental sulfur by ANME or their syntrophic partners. Other alternative electron acceptors for AOM are well documented, including nitrate, nitrite, metals, and humics [55, 139–144]. Methane oxidation linked to elemental sulfur or polysulfide reduction has been suggested for ANME-1 [145] and *Methanoperedenaceae* (formerly ANME-2d) [146] without a syntrophic bacterial partner. ANME-1b were present in our microcosms (Fig. 3B), but ANME-2c and their partner SEEP-SRB2 were more FeSRB2 did not express *phsA*, suggesting they do not reduce thiosulfate or elemental sulfur via this pathway.

However, direct thiosulfate reduction by *dsr* is possible [147–149], and both bulk *dsrA/B* transcripts and some associated with individual SEEP-SRB2 lineages were upregulated in thiosulfate-plus-methane treatments compared to thiosulfate alone (Fig. 5A, 6B). Sulfur isotopes also support this mechanism as the reduction of sulfate and thiosulfate produces fractionation of similar magnitude [21, 23]. Without methane, differences in net ε³⁴S between sulfate and thiosulfate amendments may stem from the different substrates available for *sox*-mediated oxidation. With the addition of methane and decreased relative importance of *sox*, ε^34^S in both microcosms converge around -9 ‰. Unlike thiosulfate, a mechanism for direct reduction of elemental sulfur by SEEP-SRB2 in syntrophy with ANME during AOM is not clear.

Second, AOM may also be indirectly supported by sulfite generated during *phs*-mediated thiosulfate cleavage. *PhsA* transcripts were identified in the *Desulfocapsaceae, Desulfobulbaceae*, and *Sulfurovaceae*, lineages not known to form direct partnerships with ANME. Their expression tracked sulfur source rather than methane, suggesting activity independent of syntrophy. Nevertheless, sulfite accumulated in thiosulfate-amended microcosms both with and without methane, providing a substrate that SEEP-SRB2 could reduce via *dsr*, as previously discussed for thiosulfate. Thus, *phs*-encoding microorganisms may indirectly link thiosulfate to AOM without requiring full disproportionation.

Finally, conventional sulfate-coupled AOM may be supported by the sulfate generated by disproportionation or oxidation of elemental sulfur or thiosulfate. Sulfide concentrations and sulfur isotopes may help rule out disproportionation in elemental sulfur-amended microcosms, but this process cannot be entirely excluded as a source of sulfate in the thiosulfate-amended microcosms. In both elemental sulfur- and thiosulfate-amended microcosms, *sox* transcription suggests sulfate production, though no net accumulation occurred due to rapid reduction. While some studies observed sulfate accumulation under both thiosulfate- and elemental sulfur-plus-methane conditions, which they attributed to disproportionation [41, 42], others reported no detectable sulfate in thiosulfate-plus-methane incubations and only minimal production in elemental sulfur-plus-methane incubations [43]. Discrepancies between prior reports and the current study suggest that sulfur cycling, and its link to AOM, in different seep sediments is mediated by distinct microbial taxa. This variability, evident even within ANME–SRB partnerships [1, 150], likely extends to other community members, underscoring that community composition fundamentally shapes sulfate dynamics across seep sediments.

Together, these results reveal that AOM in seep sediments may be sustained not only by direct sulfate reduction by SEEP-SRB2 but also by thiosulfate reduction or indirect pathways involving *phs*- and *sox*-dependent transformations. These results highlight that organisms such as the *Desulfocapsace*, *Desulfobulbaceae*, and *Sulfurovaceae* may be as essential to AOM as ANME-SRB. Particularly within the SMTZ, where sulfate concentrations are minimal, sulfur redox intermediates, and the taxa that contribute to their transformation, may play a central role. More broadly, this highlights the ecological and metabolic complexity underpinning methane oxidation in marine sediments.

## Conclusion

Long-term incubations of SMTZ sediments demonstrate that sulfur intermediates such as thiosulfate, sulfite, and elemental sulfur are not incidental byproducts but key components of microbial metabolism in seep environments. Methane consistently stimulated sulfide production regardless of the sulfur source, and isotopic and transcriptomic data together point toward dynamic sulfur transformations. The modest isotopic fractionations observed, combined with widespread transcription of both oxidative and reductive sulfur pathways, implicate diverse processes that may support AOM without steady sulfate buildup and underscore how cryptic cycling can mask the underlying complexity of these sediments. Viewed as a whole, these results reposition the SMTZ from a narrow zone of sulfate-coupled AOM to a more complicated biogeochemical interface where multiple microbial lineages route electrons through sulfur intermediates to sustain methane oxidation. These diverse communities likely maintain greater stability under variable redox conditions and may gain ecological advantages when sulfate supplies fluctuate. Such metabolic flexibility may become increasingly important as warming seawater influences methane release from the seafloor [151, 152]. It also reshapes how biogeochemical signals are interpreted, as the cycling of sulfur redox intermediates decouples carbon oxidation from sulfate gradients. When diffusion from the SWI is not the only source of sulfate, sulfate concentration profiles alone cannot reliably constrain remineralization rates. Accounting for this cryptic sulfur cycling is therefore crucial for interpreting sulfur isotope signatures in both modern sediments and the geological record, and for refining estimates of the global carbon budget.

## Acknowledgements

We would like to thank the captains, crew, and shipboard scientists of the R/V Western Flyer and ROV Doc Ricketts along with the Monterey Bay Aquarium Research Institute. We thank the Simons Foundation, the Gordon and Betty Moore Foundation, and the Canadian Institute for Advanced Research Fellow (CIFAR) in the Earth 4D consortium for providing funding that supported this work.

