## Supplemental Information for "Cycling of sulfur redox intermediates drives microbial activity in the sulfate–methane transition zone of cold methane seeps"

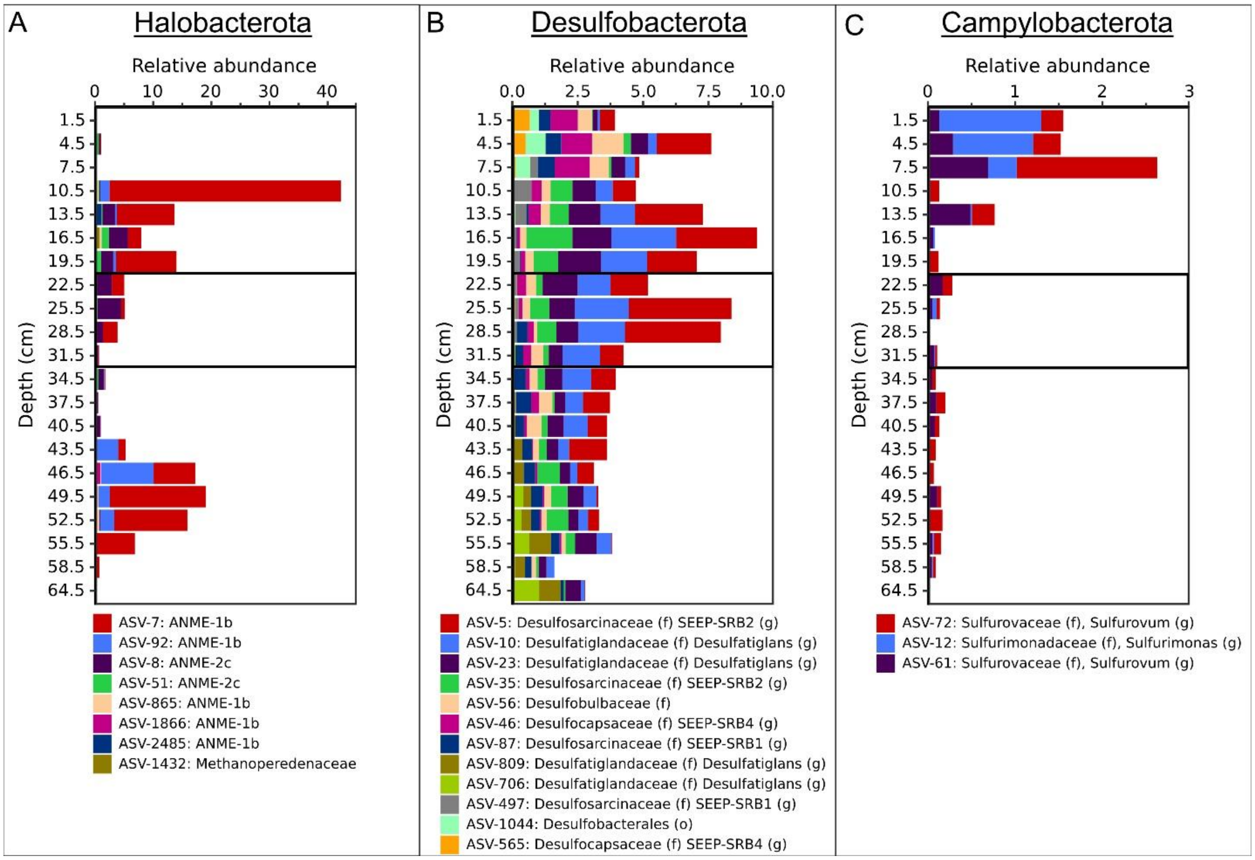


Figure S1. Community composition of seep sediment used for incubations. (A) Desulfobacterota and (B) Halobacterota ASVs > 0.6%. Highest taxonomic assignment is reported; family is reported in cases where genus level assignment was identified. The homogenized incubation horizons are outlined with a black box.

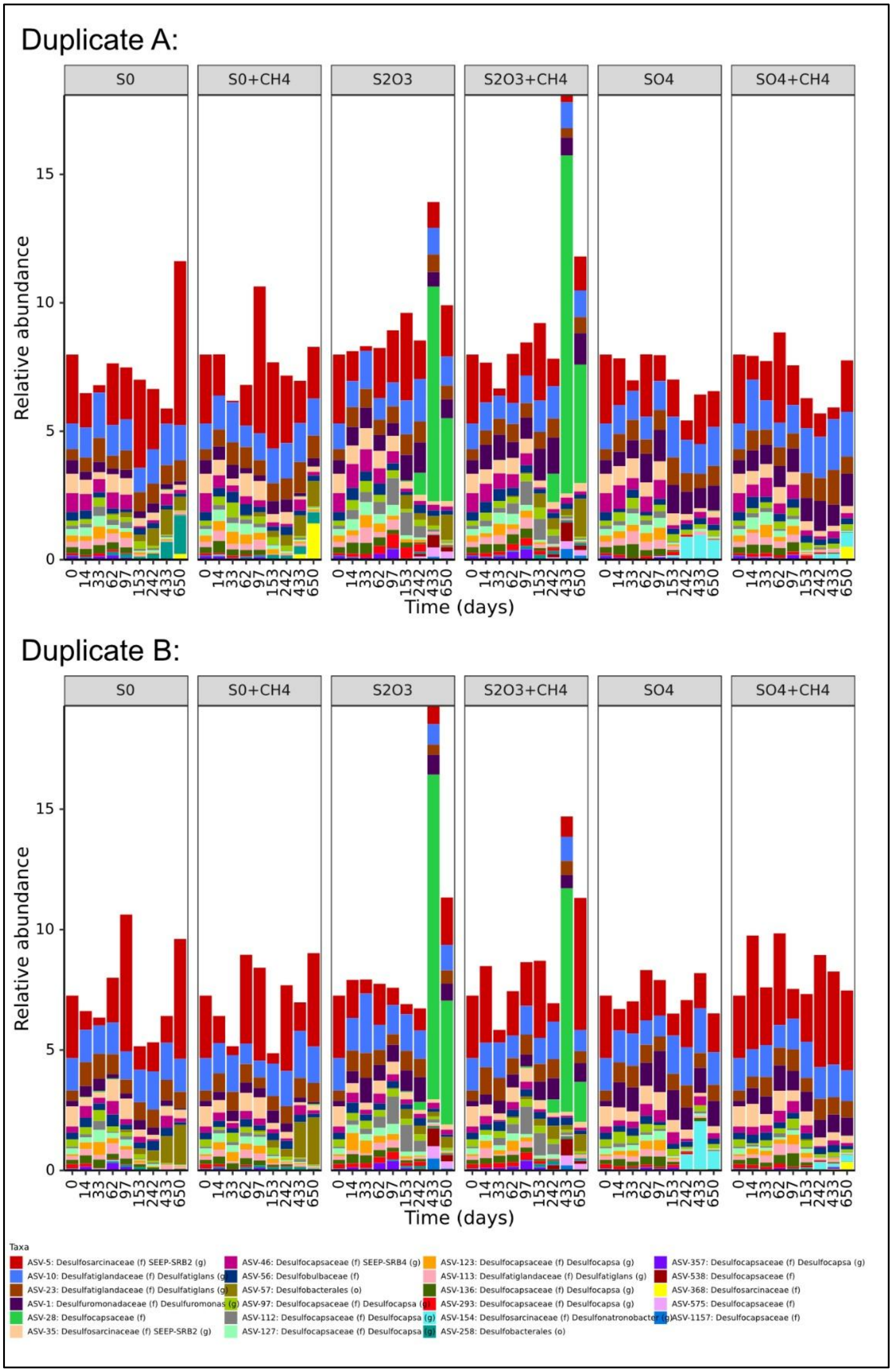


Figure S2. Temporal distribution of Desulfobacterota ASVs exceeding 0.4% relative abundance in at least two timepoints across duplicate incubations.

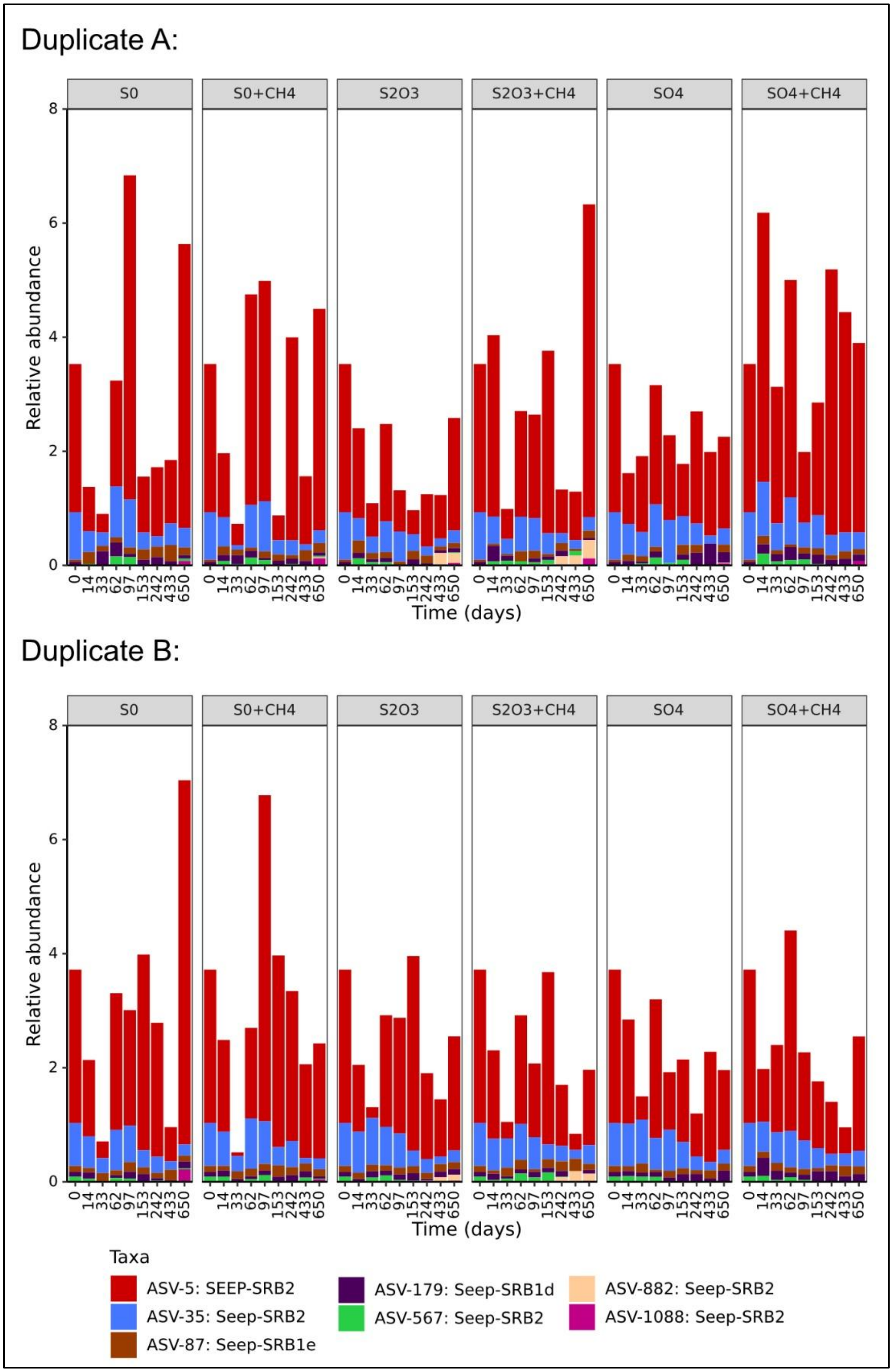


Figure S3. Temporal distribution of SEEP-SRB ASVs exceeding 0.4% relative abundance in at least two timepoints across duplicate incubations.

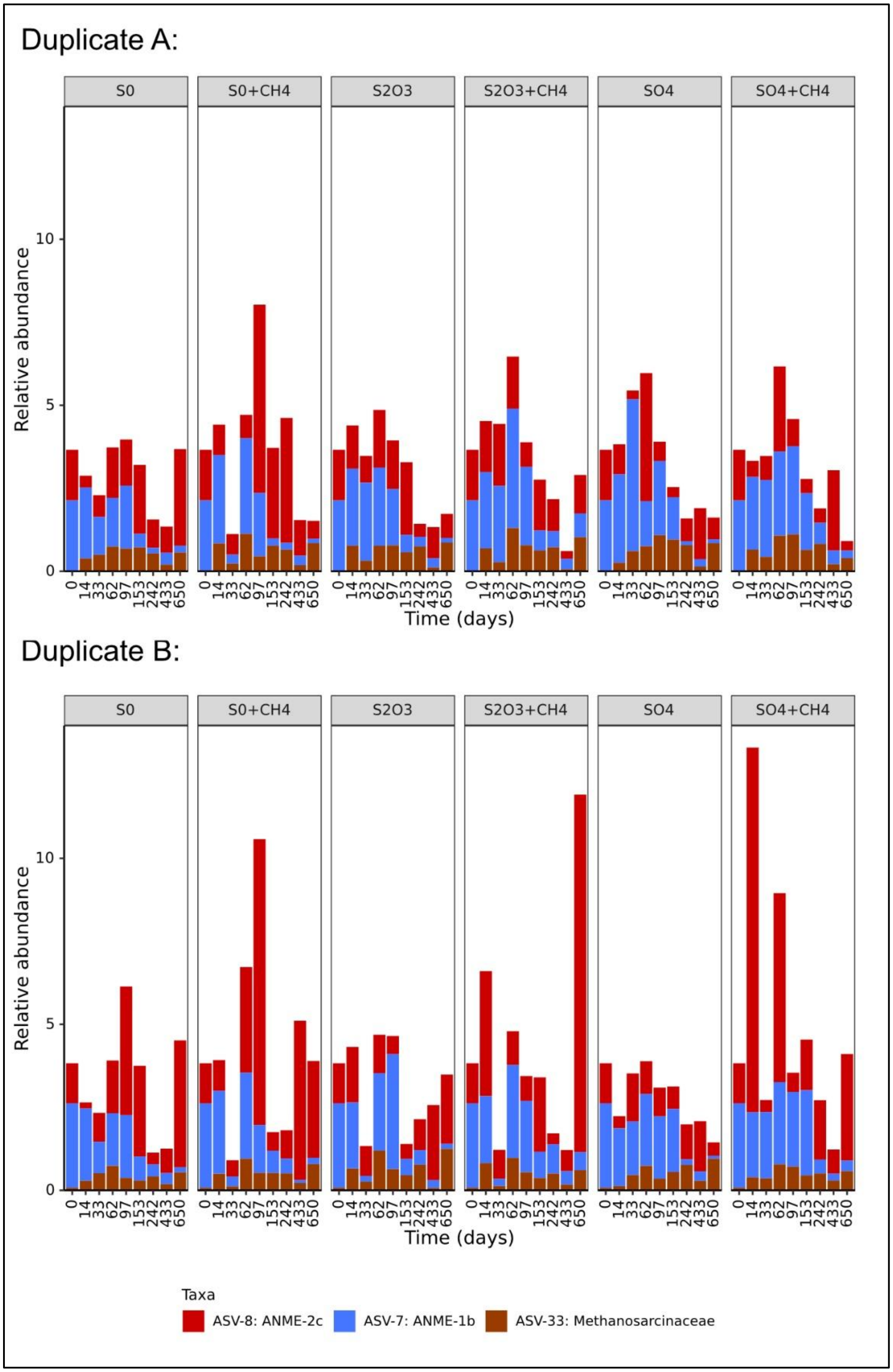


Figure S4. Temporal distribution of Halobacterota ASVs exceeding 0.4% relative abundance in at least two timepoints across duplicate incubations.

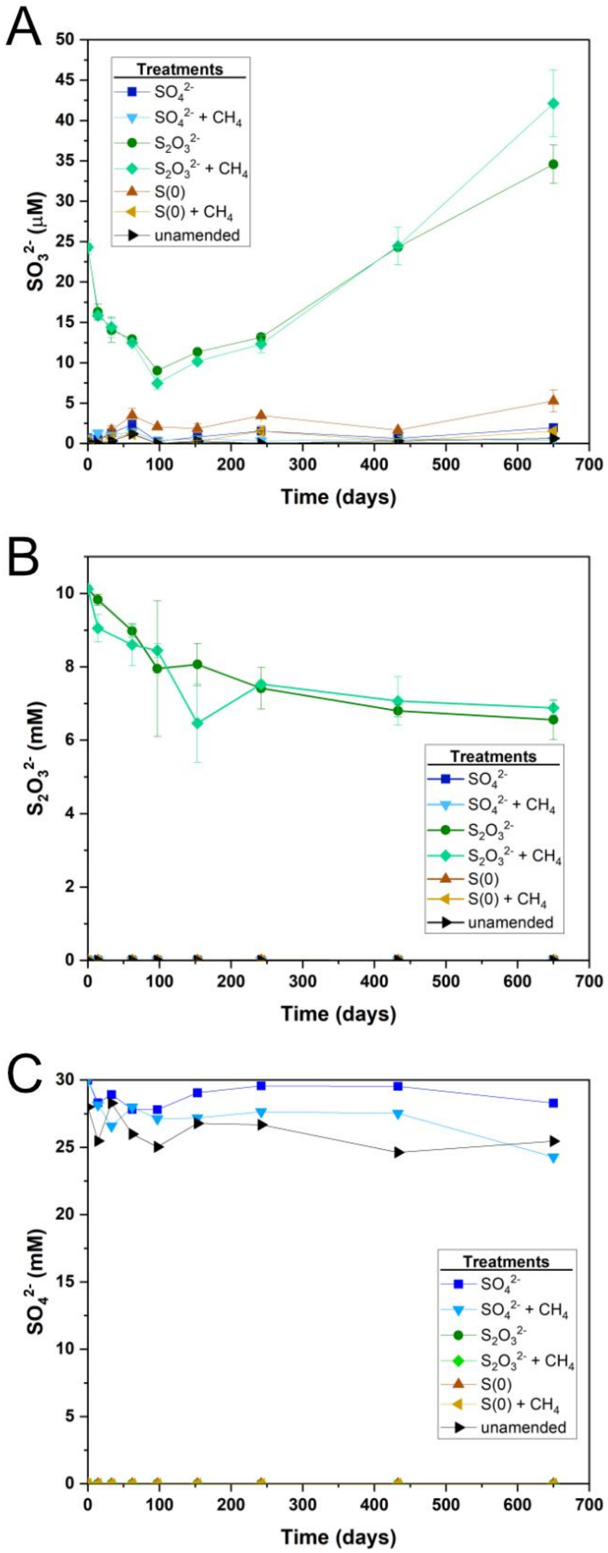


Figure S5. Concentrations of (A) sulfide, (B) thiosulfate, and (C) sulfate measured over time in anoxic microcosm incubations with different sulfur sources and methane treatments.

Figure S6. Complete phsA phylogenetic tree

Figure S7. Complete soxC phylogenetic tree

Figure S8. Complete soxY phylogenetic tree

Figure S9. Complete aprA phylogenetic tree

Figure S10. Complete dsrA phylogenetic tree

Figure S11. Complete dsrB phylogenetic tree

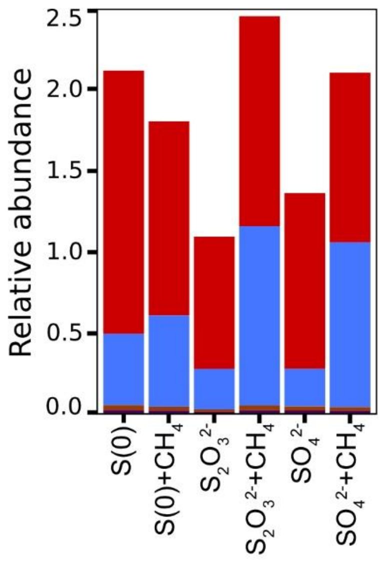


Figure S12. Relative abundances of active ANME-1 as determined from transcribed SSU rRNA.

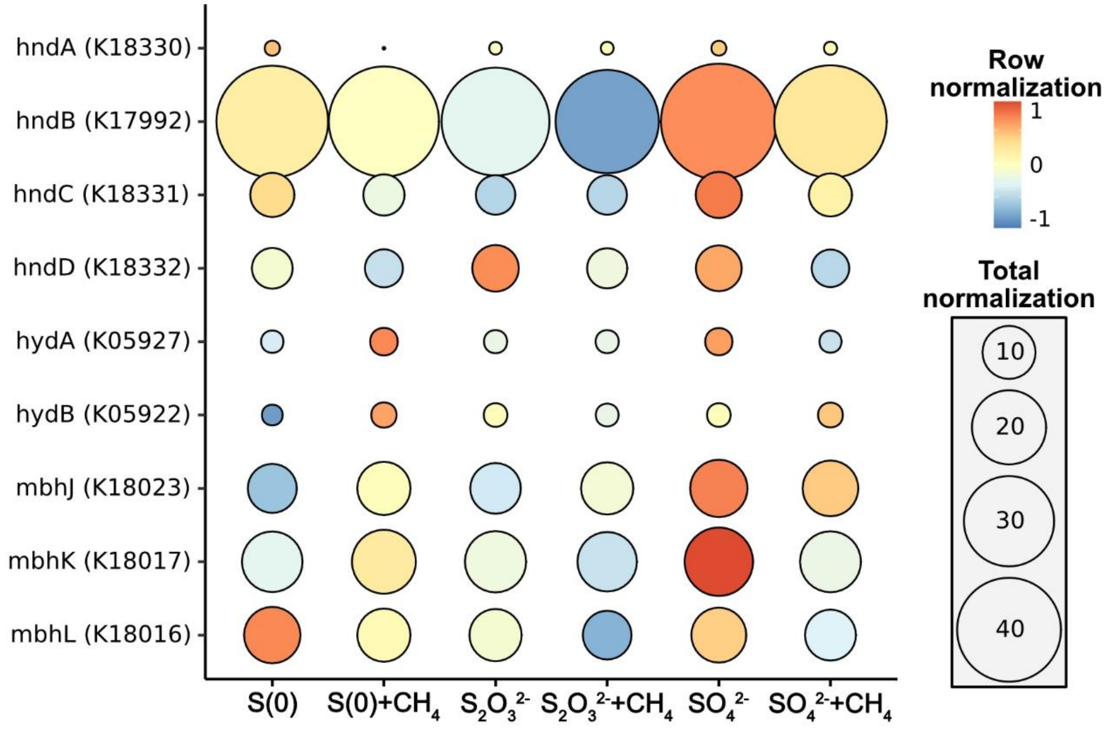


Figure S13. Transcriptomic evidence of hydrogenase activity as shown with bulk transcript abundances normalized by treatment (color) and overall transcript abundance (size).

Table S1. Sampling metadata for sediment cores collected during Monterey Canyon methane seep dives. Listed are dive numbers, push core identifiers, collection dates, geographic coordinates, water depth, seep region, and surface characteristics.

| Dive | Push Core | Date | Lat. (°N) | Long. (°W) | Depth (m) | Region | Surface Description |
|---|---|---|---|---|---|---|---|
| DR1170 | LC67 | 8/7/2019 | 36.776735 | -122.085015 | 958.6 | N Clam Bed (North Ducky) | peach mat |
| DR1170 | PC50 | 8/7/2019 | 36.77636 | -122.084905 | 966.6 | Ducky Seep | yellow mat |
| DR1170 | PC76 | 8/7/2019 | 36.77636 | -122.084905 | 966.6 | Ducky Seep | yellow mat |
| DR1170 | PC60 | 8/7/2019 | 36.77633 | -122.0848767 | 968.4 | Outside Ducky Seep | |
| DR1173 | PC41 | 8/8/2019 | 36.77639 | -122.0849517 | 965.2 | Ducky Seep | yellow mat |
| DR1171 | LC63 | 8/8/2019 | 36.77612 | -122.085585 | 960 | SW Clams 1 | yellow mat |
| DR1030 | LC46 | 5/22/2018 | 36.7760583 | -122.085615 | 956.5 | SW Clams 1 | yellow mat |

Table S2. Diversity of expressed genes involved in sulfur and methane metabolism across incubation treatments. Genes are listed with corresponding KEGG Orthology (Kofam) identifiers. Values represent the total number of unique transcripts observed across all incubations and in individual microcosms.

| | | **Number of Unique Transcripts** | | | | | | |
|---|---|---|---|---|---|---|---|---|
| **Gene** | **Kofam** | **Sum of all incubations** | **S(0)** | **S(0) + $CH_4$** | **$S_2O_3^{2-}$** | **$S_2O_3^{2-}$ + $CH_4$** | **$SO_4^{2-}$** | **$SO_4^{2-}$ + $CH_4$** |
| phsA | K08352 | 75 | 42 | 33 | 40 | 48 | 54 | 54 |
| sorA | K05301 | 6 | 3 | 2 | 5 | 4 | 2 | 4 |
| soxB | K17224 | 31 | 16 | 14 | 22 | 18 | 21 | 17 |
| soxA | K17222 | 71 | 26 | 24 | 30 | 30 | 34 | 32 |
| soxX | K17223 | 97 | 33 | 35 | 40 | 41 | 51 | 36 |
| soxY | K17226 | 170 | 70 | 54 | 74 | 68 | 93 | 74 |
| soxZ | K17227 | 130 | 46 | 41 | 49 | 46 | 74 | 56 |
| soxC | K17225 | 41 | 23 | 16 | 26 | 29 | 33 | 25 |
| soxD | K22622 | 149 | 78 | 64 | 69 | 74 | 89 | 81 |
| sqr | K17218 | 538 | 282 | 218 | 301 | 297 | 361 | 312 |
| fccA | K17230 | 336 | 140 | 103 | 136 | 150 | 180 | 157 |
| soeA | K21307 | 52 | 35 | 23 | 27 | 33 | 34 | 28 |
| aprA | K00394 | 237 | 112 | 95 | 128 | 127 | 152 | 147 |
| aprB | K00395 | 310 | 114 | 75 | 130 | 132 | 168 | 146 |
| dsrA | K11180 | 183 | 83 | 65 | 86 | 89 | 118 | 116 |
| dsrB | K11181 | 118 | 49 | 36 | 57 | 58 | 86 | 73 |
| mcrA | K00399 | 17 | 10 | 13 | 13 | 13 | 12 | 13 |
| mcrB | K00401 | 15 | 13 | 11 | 10 | 14 | 11 | 12 |
| mcrC | K03421 | 19 | 12 | 12 | 11 | 13 | 14 | 14 |
| mtrA | K00577 | 105 | 45 | 32 | 53 | 59 | 64 | 55 |
| mtrC | K00579 | 14 | 7 | 9 | 9 | 8 | 11 | 10 |
| mtrD | K00580 | 20 | 15 | 9 | 12 | 14 | 13 | 16 |
