## Supplementary figures and images for "Cycling of sulfur redox intermediates drives microbial activity in the sulfate–methane transition zone of cold methane seeps"

### Figure S6

**Figure S6. PhsA phylogenetic tree**

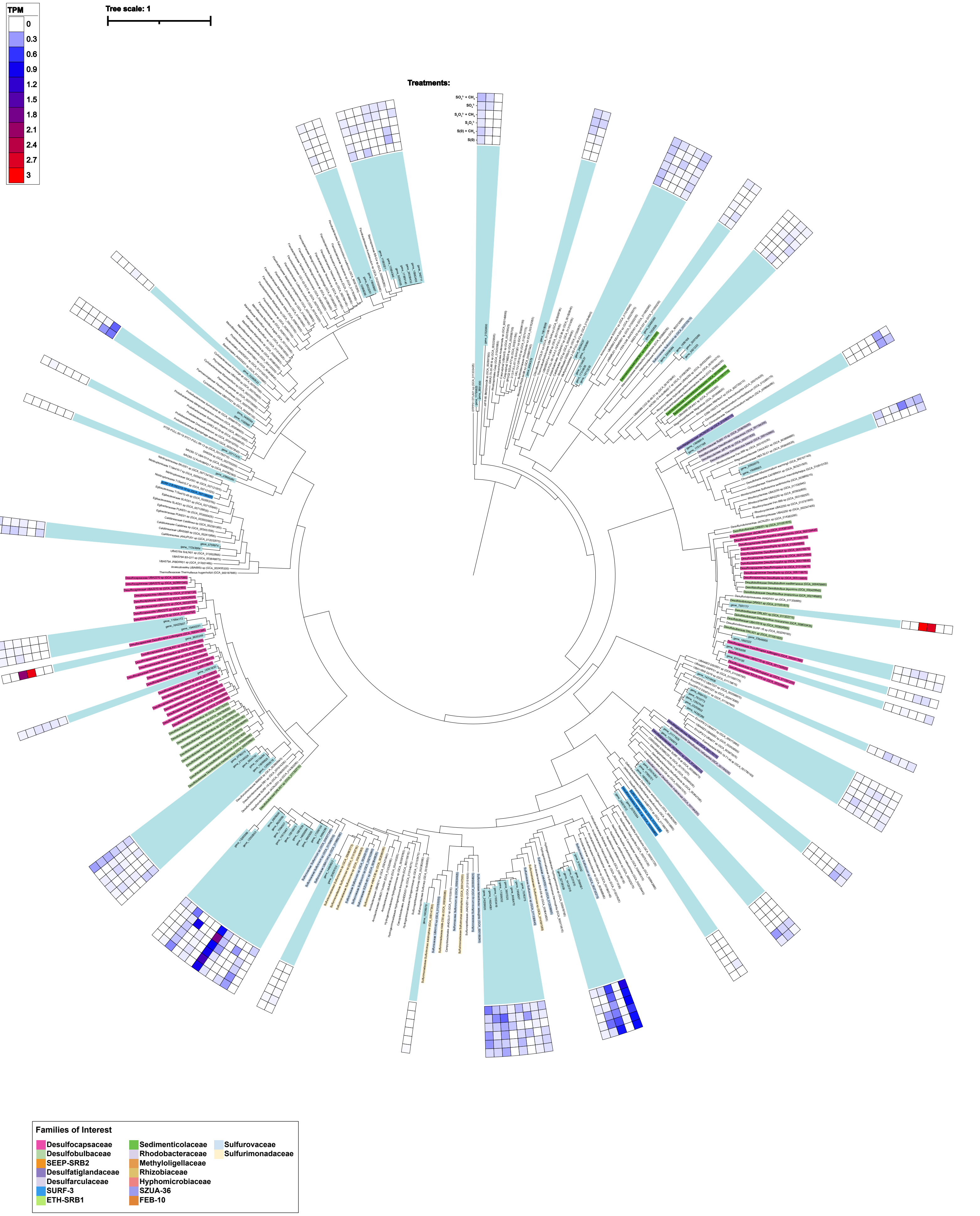

### Figure S8

**Figure S8. SoxY phylogenetic tree**

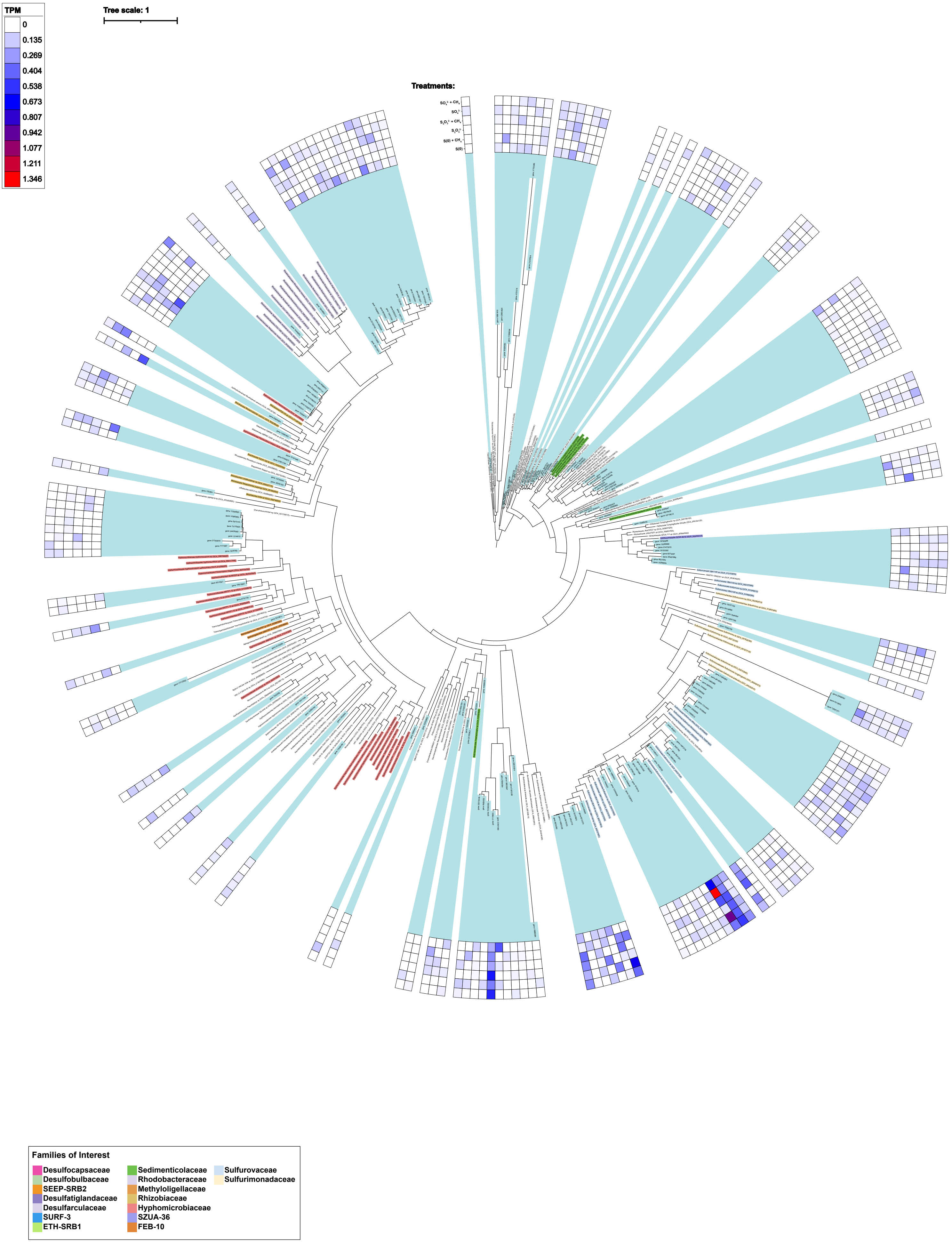

### Figure S9

**Figure S9. AprA phylogenetic tree**

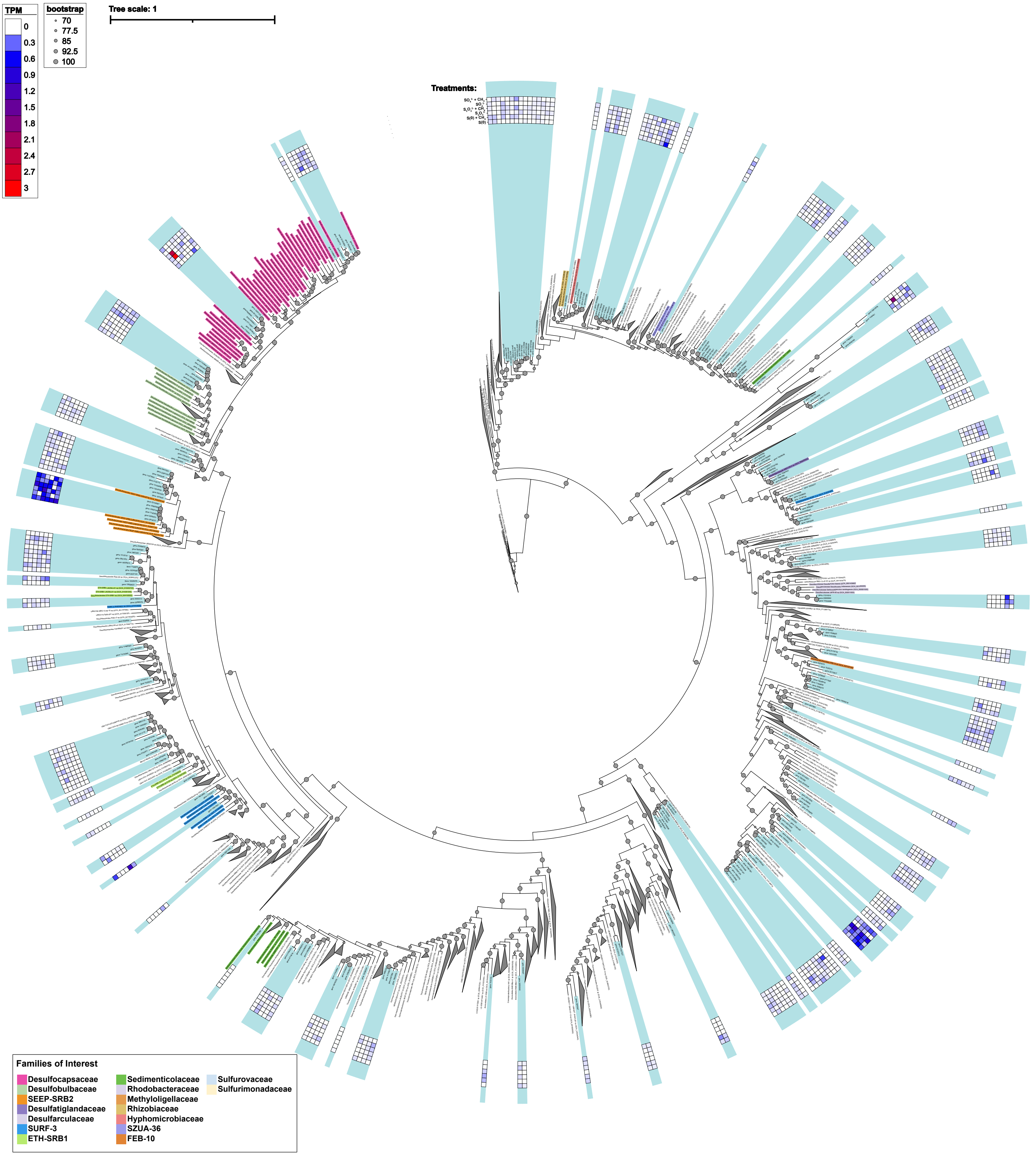

### Figure S9

**Figure S7. SoxC phylogenetic tree**

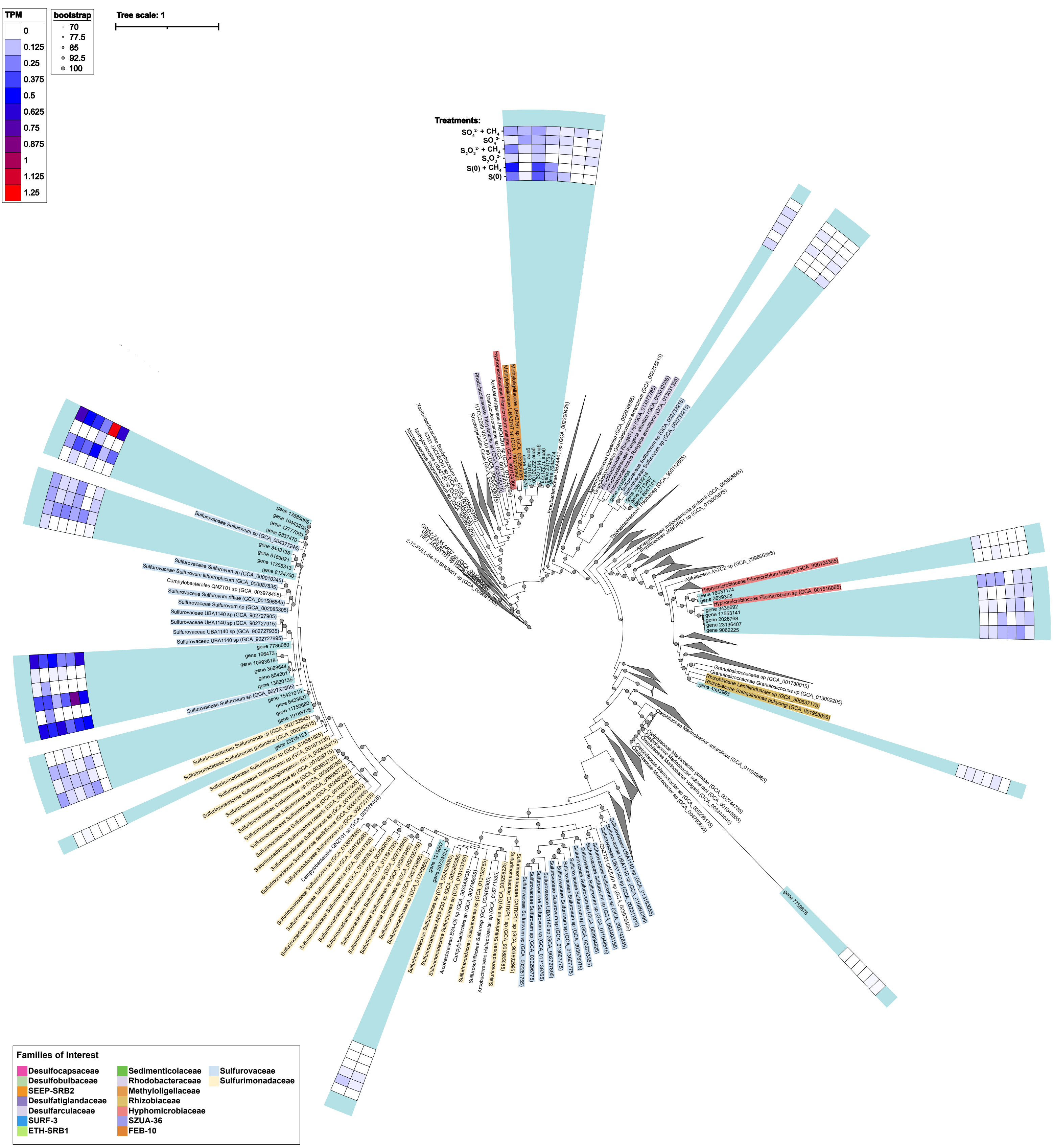

### Figure S10

**Figure S10. DsrA phylogenetic tree**

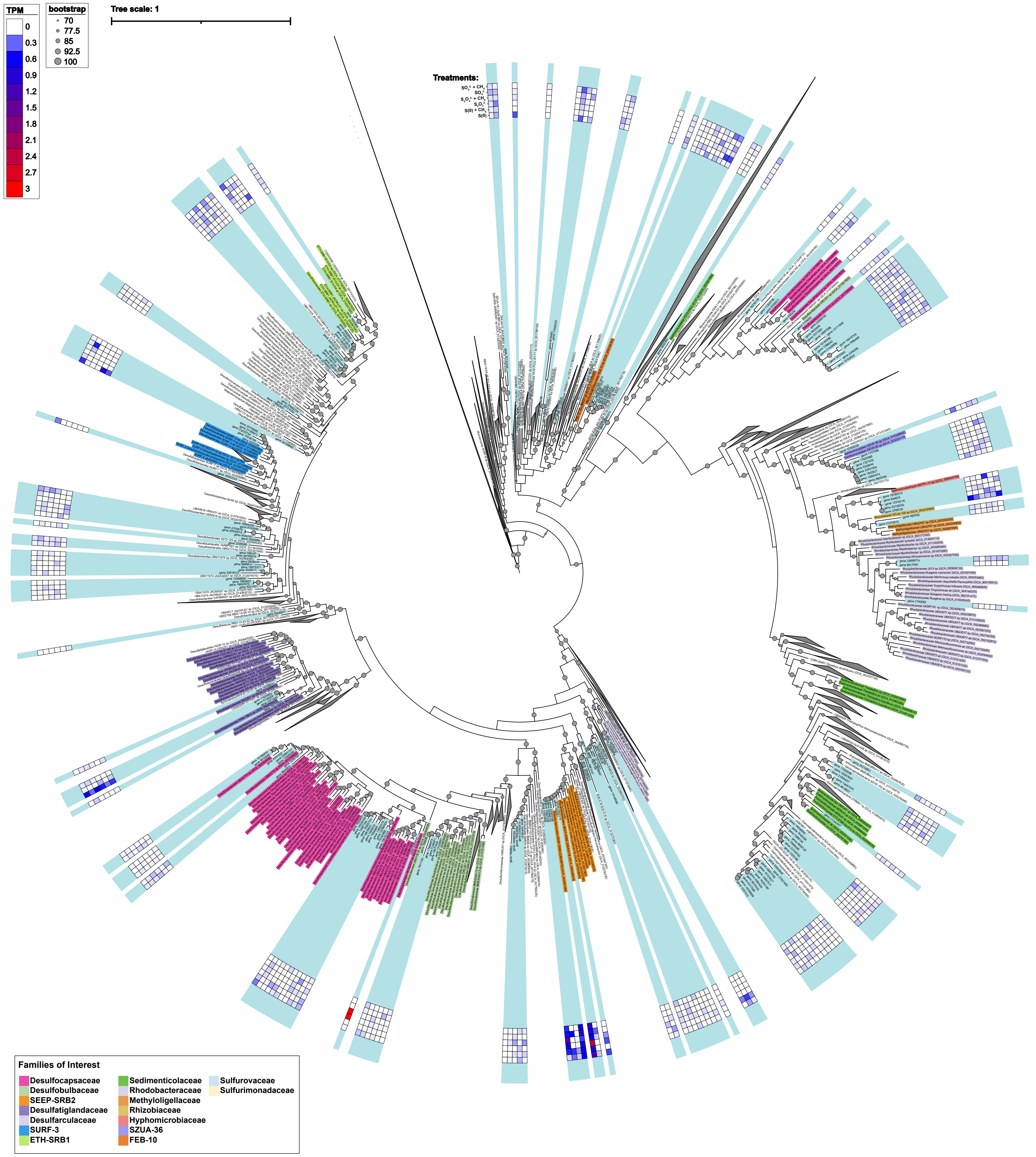

### Figure S11

**Figure S11. DsrB phylogenetic tree**

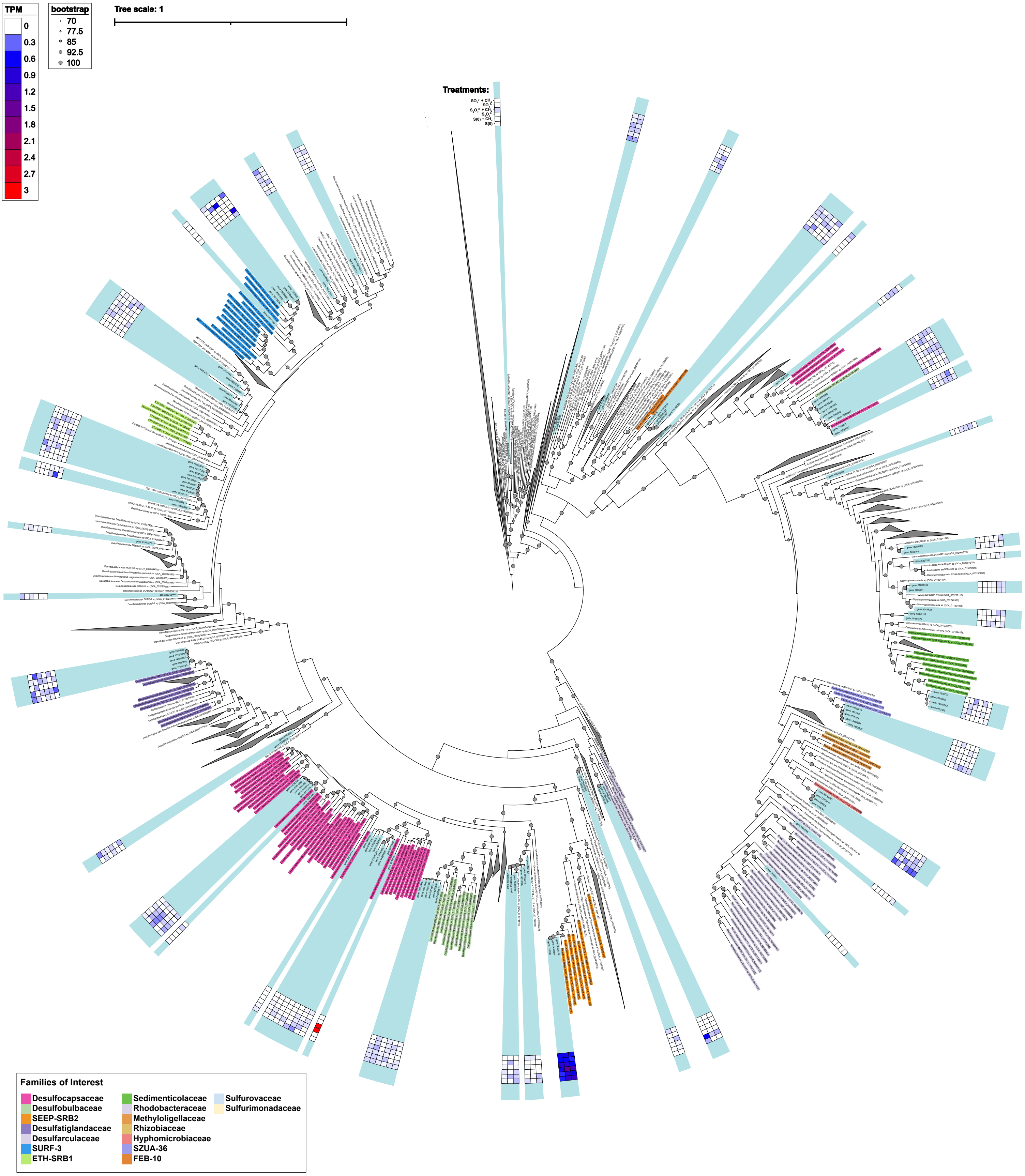
